# Spatiotemporal dynamics of β-lactam-resistant *E. coli* in young diseased calves in Wallonia, Belgium

**DOI:** 10.64898/2026.01.21.700780

**Authors:** Virginie Guérin, Nicolas Cabanel, Gan Min Marissa Diana Meijer, Guilhem Royer, Jean-Noël Duprez, Jacques G. Mainil, Marc Saulmont, Damien Thiry, Philippe Glaser

**Affiliations:** Unité EERA, Institut Pasteur, Paris France; Université Paris-Cité, Paris France; Veterinary bacteriology, Department of Infectious and Parasitic Diseases, FARAH and Faculty of Veterinary Medicine, ULiège, 4000 Liège, Belgium; Regional Animal Health and Identification Association (ARSIA), 5590 Ciney, Belgium

## Abstract

Calves are one of the most common carriers of antibiotic-resistant bacteria among farm animals. However, the impact of antibiotic usage on resistance mechanisms, transmission routes between farms, and the transmission of resistant bacteria to humans remain largely unknown. Here we analyzed the population of β-lactam resistant *E. coli* isolated over five calving seasons on 444 farms scattered throughout Wallonia, Belgium. Restrictions on critical antibiotics usage led to a reduction of resistance to 3^rd^ generation cephalosporins but has no impact on population structure and β-lactamase genes indicating a resilient population. The correlation between short genetic distances and geographic proximity suggests indirect transmission between farms by fomites with differences between regions east and west of the river Meuse. Phylogenetic analysis of calf isolates with isolates from public databases indicates transitions from bovine to human adaptation. These findings provide new means to further model the spread of *E. coli* in livestock farming.

## Introduction

*Escherichia coli* is a ubiquitous bacterium, colonizing the gut of humans and animals. It is primarily a commensal but also responsible for digestive and extra-intestinal infections, mainly urinary tract infections, sepsis and meningitis in humans and animals, or mastitis in cattle^1,2^. In young calves, *E. coli* is one of the most frequent causes of diarrhea and septicemia^3^, a major problem in their management resulting in major economic losses^4^. This leads to heavy use of antibiotics in calf farming^5^. The use of antibiotics in humans and animals contributes to the selection for antibiotic resistance and to the emergence and dissemination of multidrug-resistant (MDR) bacteria resistant to at least three classes of antibiotics^6^. The consequent increase of MDR *E. coli* and their transmission between humans, animals and the environment represent a One Health threat^7^.

In 2005, the World Health Organization published a first ranking of medically important antimicrobials, which is regularly updated^8^. This provided guidelines for restricting the use of antibiotics in food-producing animals^9^, as a major share of antibiotics is used in this sector^10^. In Belgium, a Royal Decree applied since 2016 restricts the use of antibiotics considered critical to human health like 3^rd^ and 4^th^ generation cephalosporins (3GC and 4GC) in production animals^11^. To analyze the impact of this regulation, the β-lactam resistance profiles of 3917 *E. coli* isolates from 3537 calves were analyzed^12^. These were collected during seven calving seasons (2014 to 2021) in Wallonia, Belgium. Analysis of the trends between the seven calving seasons showed a decrease of the two β-lactam resistance profiles associated with 3GC resistance, namely extended-spectrum β-lactamase (ESBL) and cephalosporinase (AmpC). This reduction of resistance to 3GC was compensated by an increase of isolates resistant to amoxycillin but not to 3GC and was likely due to the decreased use of 3GC in farm animals^12^. The resistance rate among a bacterial species results from a combination of selection and transmission^13^. Transmission routes are difficult to track and require combining epidemiological and genomic data^14^. The transmission of isolates between animals and humans by whole genome sequencing (WGS) has been addressed at the national level by genomic analyses of bloodstream-associated *E. coli* from the United Kingdom, and isolates from livestock farms^15^ and of colistin-resistant Enterobacterales in humans, livestock farms and the environment in Belgium and The Netherlands^16^. These studies suggested rare cases of transmission between animals and humans. On the other hand, a systematic analysis of ESBL *E. coli* in humans, poultry and food in a more limited area in Antananarivo showed substantial circulation of strains and plasmids within and between these three sectors^17^. However, spatiotemporal dynamics of drug-resistant *E. coli* at a country level remains largely uncharacterized.

Through the surveillance of β-lactam resistant *E. coli* infecting calves we have obtained a dense sampling of isolates from Wallonia. Here, we characterize the temporal and geographic dynamics of *E. coli* in sick calves by sequencing 764 isolates from 444 farms dispersed throughout Wallonia during five calving seasons (2014 to 2020). Despite the reduction in the rate of 3GC resistance, we did not observe a significant modification of the population structure and β-lactam resistance mechanisms, suggesting a resilience of the population. By correlating genetic and geographical data we quantified the rate of dissemination and observed different patterns depending on the different breeding regions. Furthermore, by including genome sequences from the international database EnteroBase^18^ in the phylogenetic analysis, we showed that locally disseminated lineages of calf isolates include isolates that infect humans, suggesting their zoonotic potential.

## Results

### Diversity of *E. coli* from diseased calves isolated in Wallonia

We selected 681 isolates for WGS from 1,495 ampicillin-resistant *E. coli* collected from 831 farms over three calving seasons (S1 to S3 November to February 2017 to 2020) (Table S1)^12^. Isolates were chosen to keep the balance between the four β-lactam resistance profiles across the three calving seasons (see Methods section, Table 1). They were from 379 farms spread out over Wallonia. We first evaluated their diversity by performing whole genome phylogeny including 83 isolates from the two previous calving seasons (Fig. 1). Despite their isolation from diseased animals, these isolates belong to all phylogroups (A, B1, C, D, E, F and G)^19^ except B2 associated with extraintestinal infections in humans^1^.

**Fig. 1.**
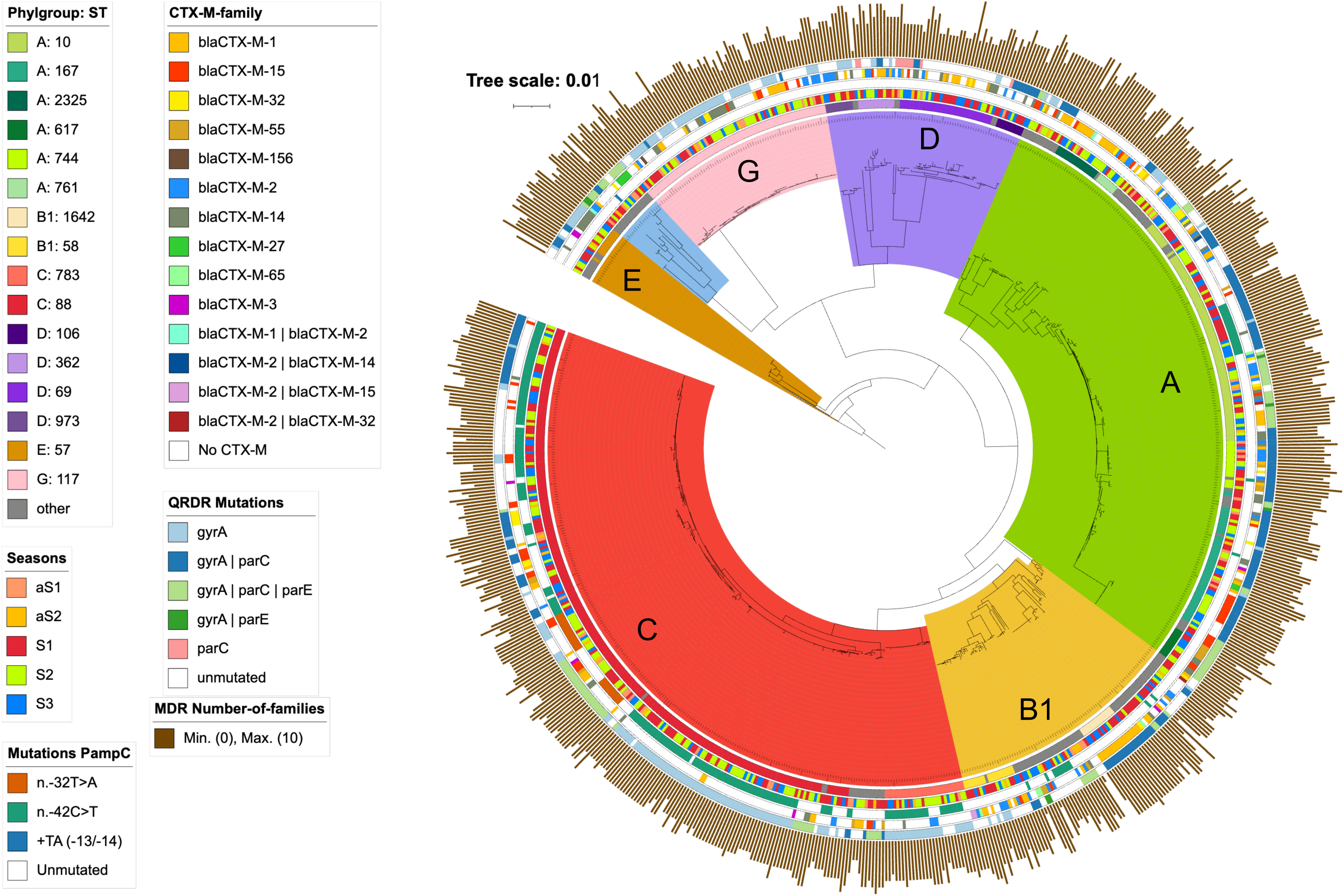
Phylogeny and characteristics of the 764 *E. coli* isolates from calves. Phylogeny was performed by maximum likelihood by using Roary, 3.13.0 ^46^ and RAxML 8.2.12, ^47^ with *Escherichia fergusonii* (NZ_CP057657) as outgroup. Genomes were annotated according to the figure key as circles from inside to outside: the main ST and the corresponding phylogroup, the calving season, P*_ampC_* mutations, *bla*_CTX-M_ genes, mutations in the *gyrA* and *parC* quinolone resistance determining regions and *parE*, and the number of resistances to antibiotic families based on genes and mutations identified. Details are in Table S1 and S2. Phylogroups are indicated by colored sectors. The tree was visualized with iTOL v7.^49^

**Table 1:** Collected and sequenced *E. coli* isolates according to the calving seasons and the β-lactam resistance profile.

| & β-lactam<br>resistance<br>profile | Pathogenic |  |  |  |  |  | Non-pathogenic |  |  |  |  |  | Total<br>WGS/collected |
| --- | --- | --- | --- | --- | --- | --- | --- | --- | --- | --- | --- | --- | --- |
|  | S1 (2017-2018) |  | S2 (2018-2019) |  | S3 (2019-2020) |  | S1 (2017-2018) |  | S2 (2018-2019) |  | S3 (2019-2020) |  |  |
|  | §WGS | Collected | WGS | Collected | WGS | Collected | WGS | Collected | WGS | Collected | WGS | Collected |  |
| NSBL | 47 | 327 | 37 | 277 | 39 | 208 | 0 | 8 | 3 | 11 | 1 | 7 | 127/838 |
| ESBL | 61 | 64 | 69 | 72 | 32 | 48 | 72 | 91 | 60 | 75 | 35 | 38 | 329/388 |
| AmpC | 9 | 9 | 4 | 4 | 1 | 1 | 13 | 15 | 6 | 6 | 15 | 15 | 48/50 |
| AmpC-like | 40 | 45 | 36 | 42 | 23 | 29 | 31 | 36 | 29 | 37 | 18 | 22 | 177/211 |
| Susceptible | 0 | 7 | 0 | 0 | 0 | 0 | 0 | 1 | 0 | 0 | 0 | 0 | 0/8 |
| Total | 157 | 452 | 146 | 395 | 95 | 286 | 116 | 151 | 98 | 129 | 69 | 82 | 681/1495 |

<sup>&</sup> β-lactam resistance profiles (narrow spectrum β-lactamase, extended spectrum β-lactamase, AmpC, and AmpC-like, which corresponded mainly to *E. coli* chromosomal *ampC* derepression) were as previously described <sup>37</sup>. <sup>§</sup>Illumina whole genome sequenced.

The 681 isolates belong to 77 different STs, with 78% of the isolates belonging to the 13 most abundant (Fig. 1 and 2a-b). Phylogroup C was dominant with 32% of the isolates, including 205 ST88 isolates and 25 from the double locus variant, ST783. Phylogroup A contains 26% of the isolates, with 80% belonging to clonal complex 10 (76 ST10, 39 ST167 and 11 ST617). The two most common STs in phylogroup B1 were ST58 (n=17, 22%) and ST1642 (n=14, 18%). 34 of the 70 Phylogroup D isolates belong to ST69 (49%) and nine to its single variant ST106 (13%). All phylogroup G isolates belong to ST117 (n=57). Despite the decreased 3GC resistance rate previously reported^12^, the ST distribution of the isolates according to the four phenotypic categories did not change during the three seasons (Fig. 2 a-b).

**Fig. 2.**
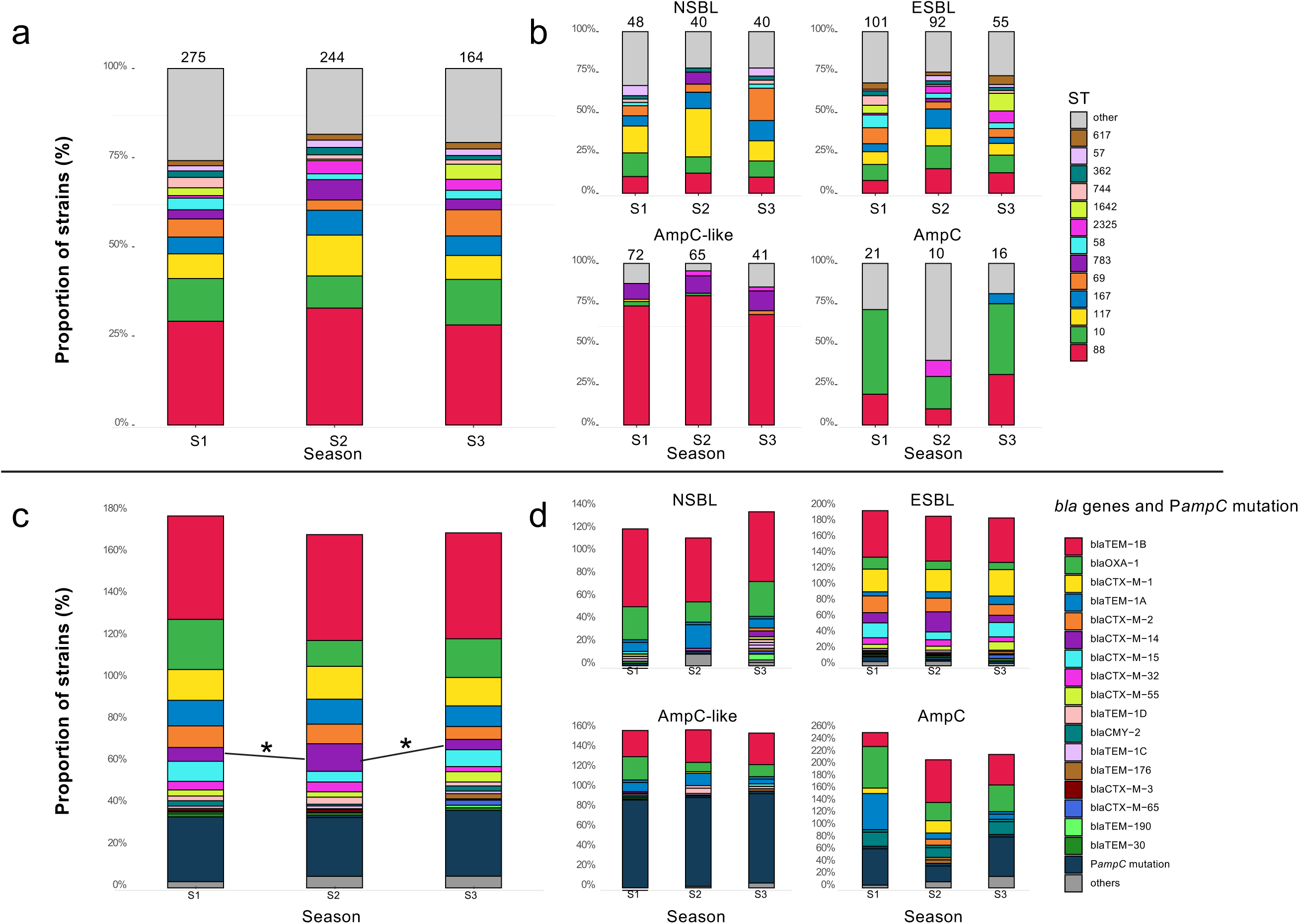
Sequence type and β-lactamase gene distribution according to season and ß-lactam resistance profile. **a**. Sequence type (ST) distribution of the 681 isolates according to the three calving seasons S1, S2 and S3. **b**. ST distribution according to the four phenotypic profiles: NSBL, ESBL, AmpC-like and AmpC. Number of isolates of each profile are indicated above the histogram. **c**. β-lactamase gene and P*_ampC_* mutation frequencies in the 681 isolates according to the three calving seasons. **d**. ß-lactamase gene and P*_ampC_* mutation frequencies according to the four ß-lactam resistance profiles: NSBL, ESBL, AmpC-like and AmpC. Percentages correspond to the percentage of isolates carrying the β-lactamase gene or the P*_ampC_* mutation. It explains why total numbers are higher than 100%. Statistical significance between β-lactamase gene and ST frequencies were assed with a global Kruskal-Wallis test. * P.<0.05. No trend was identified over the three calving seasons. ST (A and B) and ß-lactamase genes and P*_ampC_* mutation (C and D) are colored according to the figure key.

### *E. coli* isolates from seasons S1-3 showed diverse but stable patterns of β-lactamase genes

The number of acquired β-lactamase genes among isolates from seasons S1-3 ranged from none in 79 isolates, to four in two isolates (Table S1). 206 isolates (30%) had a mutation in the chromosomal P*_ampC_* promoter at position −42 or −32 or a TA insertion between positions −13 and −14. These mutations were shown to derepress *ampC* expression^20,21^.

The *bla* gene content combined with P*_ampC_* mutations was consistent with the β-lactam resistance profile (Fig. 2 c-d, Table S1). Among the 127 NSBL-producing isolates, *bla*_TEM-1_ variants and *bla*_OXA-1_ were predominant, present in 88% and 28% of the isolates respectively. All except two of the 329 isolates showing an ESBL phenotypic profile carried a *bla*_CTX-M_ gene (Fig. 2 c-d, Table S1). Six isolates carrying this ESBL gene showed an AmpC (n=4) or NSBL (n=2) profile. *bla*_CTX-M-1_ was the most frequent gene (30%), followed by *bla*_CTX-M-2_ (19%), and *bla*_CTX-M-14_ and *bla*_CTX-M-15_ (17% each). The AmpC-like phenotype (n=177) was mainly caused by mutations in the promoter region of P*_ampC_*, which were found in 90% of the isolates (n=170). Four AmpC-like isolates harbored an acquired AmpC-encoding gene, *bla*_CMY_2_ (n=1) or *bla*_DHA_1_ (n=3). Among the 48 isolates with an AmpC phenotype, 16 harbored an AmpC-encoding gene: *bla*_CMY-2_ (n=13), *bla*_DHA-1_ (n=2) or both (n=1); 28 isolates had mutations in P*_ampC_* with 26 harboring acquired β-lactamase genes, predominantly *bla*_OXA-1_ (n=18); four carried *bla*_CTX-M-1_ combined with one or two other β-lactamase genes (Table S1).

In addition to β-lactamase genes, isolates carried a wide variety of ARGs, with a median of 10, ranging from none in one isolate up to 21 in two isolates. We identified mutations in the quinolone resistance-determining regions (QRDR) of *gyrA* and/or *parC* in 67.4% of the isolates (n=459). The median number of ARGs ranged from eight for AmpC-like isolates to 14 for those with an AmpC phenotype (Fig. S2). There was no carbapenemase gene, in agreement with the absence of carbapenem-resistant isolates. On the other hand, 54 isolates from 18 different STs carried an *mcr* gene, which provides resistance against colistin. The vast majority expressed *mcr-1* (n=50), two expressed *mcr-2* and two *mcr-9*. *mcr*-carrying isolates carried a larger number of ARGs (median=13) than non-*mcr*-carrying isolates (median=10) (p<0.0001).

Despite the modification of antibiotic usage, no statistically significant trend was observed during the three calving seasons studied for the mean ARG number, β-lactamase gene diversity and number of P*_ampC_* mutations when considering the four different β-lactam resistance patterns (Fig. 2 c-d).

### Mutations in the *ampC* promoter were predominant among phylogroup C isolates

Among the 206 isolates of seasons S1-3 showing a mutation in P*_ampC_*, 177 (86%) belonged to phylogroup C (157 to ST88 and 20 to ST783). Symmetrically, P*_ampC_* mutated isolates represented 76% and 77% of the isolates from these two STs, respectively. During the two previous calving seasons, 17 out of 83 isolates (20%) showed mutation in P*_ampC_*. All belonged to phylogroup C, representing 12 of the 19 ST88 isolates (63%), three of the four ST783 isolates and the three ST23 isolates. To contextualize the acquisition of these mutations, we performed whole genome phylogeny by including genome sequences retrieved from EnteroBase: 1243 ST88 (Fig. 3) and 14 ST783 (Fig. S3b). Among them, 222 ST88 (18%) and 4 ST783 (29%) isolates showed a mutation in P*_ampC_*. Among the ST88 isolates from Enterobase, P*_ampC_* mutations were more frequently associated with livestock (31% of the 426 isolates) than with humans (12% of the 268 isolates) (p<0.001). The 24 isolates from the study mutated at position −32 (T>A) were monophyletic and belong to ST88. This lineage (L-ST88-1) includes 14 isolates from EnteroBase (Fig. 3, Table 2). Two large lineages, likely monophyletic, L-ST88-2 (74 isolates) and L-ST88-3 (63 isolates), share the same mutation at −42 (C>T) in P*_ampC_* (Fig. 3, Table 2). 24 additional ST88 isolates clustered in seven lineages of one to seven isolates showed this mutation at position −42. Among the 42 ST783 isolates, 26, including 23 from our study, share the −42 C>T mutation in P*_ampC_*. Based on the tree structure (Fig. S3b) several independent mutation events in P*_ampC_* were predicted. Among other phylogroups, we detected two additional monophyletic lineages of 18 ST10 (Fig. S3a) and six ST345 (Fig. 1) isolates. One ST624 isolate, classified as AmpC-like, carried a TA insertion between positions −13 and −14 of P*_ampC_*. We analyzed polymorphisms in regions 10 kb upstream and downstream of *ampC* to determine if the P*_ampC_* mutation was acquired by recombination. The cluster of 18 ST10 isolates mutated in P*_ampC_*, showed only two SNPs compared to the L-ST88-2 isolate R0092. The acquisition of the mutation in P*_ampC_* resulted therefore of a recombination event from a mutated ST88 strain. We did not predict any other case of P*_ampC_* mutation acquisition by recombination (Fig. S4).

**Fig. 3.**
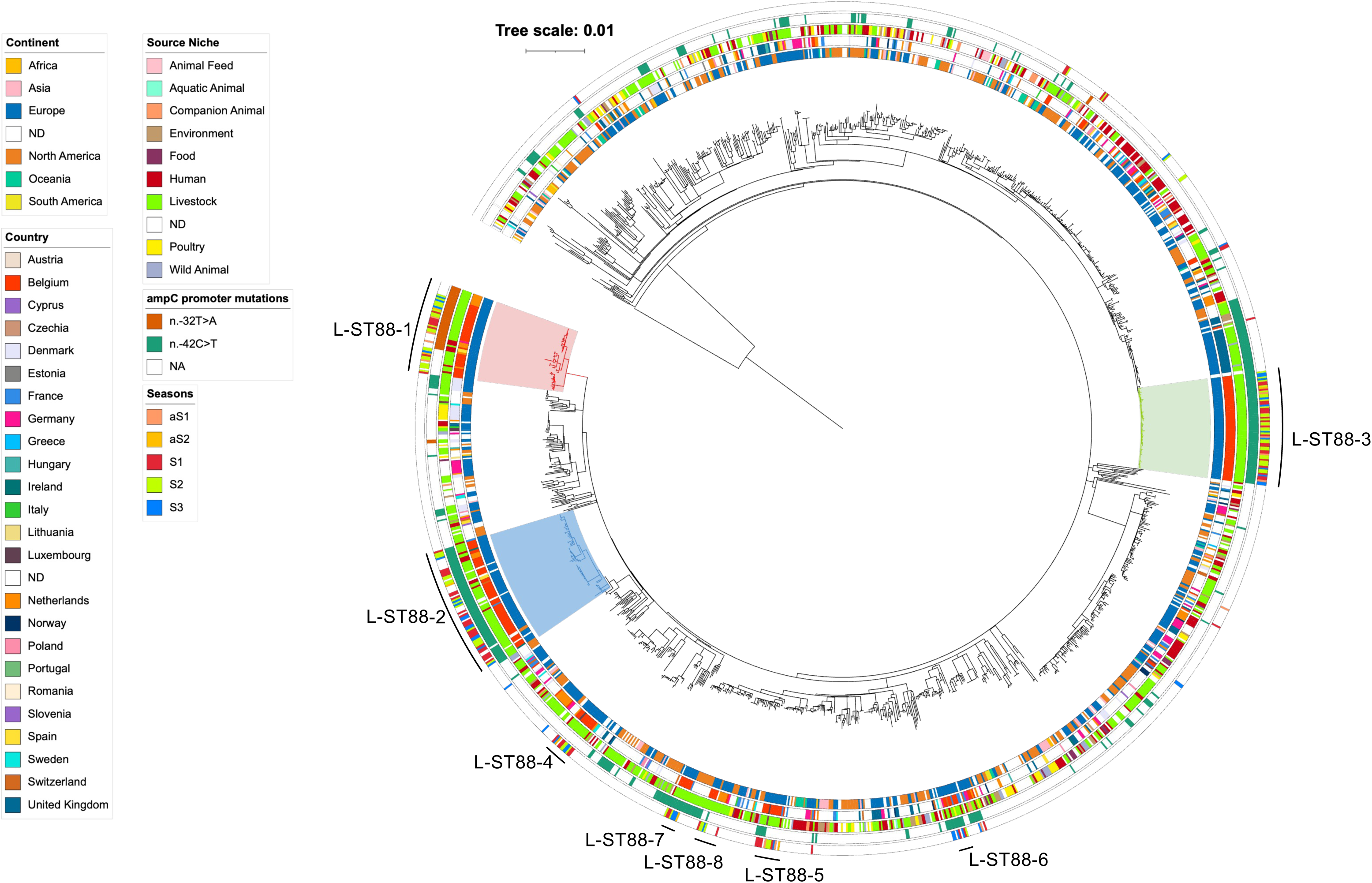
Phylogeny and characteristics of 189 ST88 *E. coli* isolates from the five seasons combined with 1243 Enterobase isolates. Maximum likelihood phylogeny was performed by using RAxML 8.2.12 ^47^ based on recombination-free core genome alignment, computed with Snippy 4.6.0 (https://github.com/tseemann/snippy) and Gubbins 2.4.1 ^48^ to filter recombined regions. Genomes were annotated according to the figure key as circles from inside to outside: the continent of origin, the country, the source niche, the P*_ampC_* mutation and visualized with iTOL v7. ^49^ The three main lineages of calf isolates mutated in P*_ampC_* are colored. Clusters reported in table 2 are indicated outside the last circle.

**Table 2:** Monophyletic lineages gathering ^&^deduplicated isolates from the study.

| ST | Lineage | # Isolates | # Study isolates | §Origin of Enterobase isolates | # Human isolates |
| --- | --- | --- | --- | --- | --- |
| **ST10** | L-ST10-1 | 12 | 10 | UK (1, farm) Italy (1, Bovine) | - |
|  | L-ST10-2 | 15 | 15 | - | - |
|  | L-ST10-3 | 17 | 17 | - | - |
|  | L-ST10-4 | 10 | 10 | - | - |
| **ST57** | L-ST57-1 | 6 | 5 | Germany (1, bovine) | - |
|  | L-ST57-2 | 8 | 5 | Germany (1, animal; 1, human), France (1, Human) | 2 |
| **ST88** | L-ST88-1 | 52 | 36 | Netherlands (7, bovine; 5 human), Spain (2, human), Belgium (1 human), Germany (1 human) | 9 |
|  | L-ST88-2 | 69 | 47 | USA (3, bovine), Ireland (1, human; 1, bovine), France (1, human), Belgium (1, Human; 2, swine; 4, bovine), Switzerland (1, human), Spain (2, human), Netherlands (3, human), Unknown (2) | 9 |
|  | L-ST88-3 | 63 | 63 | - | - |
|  | L-ST88-4 | 14 | 10 | Germany (2, human), Luxembourg (1, bovine), Netherlands (1, human) | 3 |
|  | L-ST88-5 | 13 | 11 | France (1, human), Luxembourg (1, food) | 1 |
|  | L-ST88-6 | 14 | 7 | France (2, bovine; 1, human; 1, poultry), Switzerland (2, human), UK (1, human) | 4 |
|  | L-ST88-7 | 5 | 5 | - | - |
|  | L-ST88-8 | 5 | 5 | - | - |
| **ST106** | L-ST106-1 | 7 | 7 | - | - |
| **ST117** | L-ST117-1 | 50 | 44 | France (1, human; 1, bovine), Netherlands (1, human), Luxembourg (1, swine), Sweden (1, human), USA (1, bovine) | 3 |
|  | L-ST117-2 | 15 | 14 | UK (1, Human) | 1 |
|  | L-ST117-3 | 6 | 5 | North America (1, human) | 1 |
| **ST167** | L-ST167-1 | 15 | 14 | 1 (unknown) | - |
|  | L-ST167-2 | 8 | 8 | - | - |
|  | L-ST167-3 | 12 | 8 | Germany (2, human), Netherlands (1, human), UK (1, Human) | 4 |
| **ST345** | L-ST345-1 | 6 | 6 | - | - |
| **ST457** | L-ST457-1 | 8 | 5 | Luxembourg (3, Livestock) | - |
| **ST617** | L-ST617-1 | 14 | 10 | Germany (3, Human) Luxembourg (1) | 3 |
| **ST783** | L-ST783-1 | 32 | 27 | France (2, bovine), Poland (1, bovine) Canada (1, bovine), Luxembourg (1, food) | - |
| **ST1642** | L-ST1642-1 | 13 | 13 | - | - |
| **ST2325** | L-ST2325-1 | 8 | 8 | - | - |
|  | L-ST2325-2 | 5 | 5 | - | - |
| **Number of isolates** |  | 502 | 420 | 82 | 40 |

&Only one isolates was considered for clonal isolates from the same animal. #Number of isolates. §Country or origin, in parenthesis the number of isolates and the source.

### A majority of *E. coli* isolates cluster in monophyletic lineages frequently also encompassing human isolates

To further characterize relationships between *E. coli* isolates from different herds, we analyzed phylogenies of the 13 STs with more than ten isolates, contextualized by including EnteroBase genomes (Fig. 3, Fig. S3,5,6,7). Visual inspection of phylogenetic trees showed that many calf isolates formed distinct lineages with few EnteroBase isolates. Using a threshold of five isolates from the study, we identified 28 lineages from 12 STs, comprising 420 (54%) non-duplicated isolates from the study (Table 2). Twelve lineages included only isolates (n=5 to 63) from the study. In the other 16 lineages, 82 isolates from EnteroBase were included. Only six isolates (7%) came from outside Europe (North America). Eight isolates were from Belgium, and most other isolates were from countries bordering Belgium (The Netherlands, 18; France, 10; Germany, 11; Luxembourg, 5), and from the UK (n=5) (Table 2). Thus, these lineages showed a limited geographic diffusion. 49% of EnteroBase isolates were of human origin (n=40), suggesting a transition from bovine to human host. This was confirmed by PastML^22^, predicting cattle as the most likely host of the ancestral strain of ten of the 11 lineages containing human isolates and 18 bovine host to human host transitions with a probability higher than 0.8. All EnteroBase isolates, as those from this study, were predicted as MDR and carried a median of nine ARG per strain, 51% had a P*_ampC_* mutation and 83% had *gyrA*/*parC* QRDR mutations (Table S3). These findings suggest a zoonotic potential for these MDR lineages originating from cattle.

### Dissemination occurred mainly at short distances with differences between eastern and western parts of Wallonia

In complement to the phylogenetic analysis, we computed the cgMLST distances between all pairs of isolates, after deduplicating identical isolates from the same animal (Fig. 4). It revealed a peak centered at a cgMLST distance of ten different alleles. 511 isolates had a cgMLST distance of less than 21 alleles with at least one other isolate (Fig. 4b, Table S4). Among these isolates, 354 belonged to the 28 lineages identified on the phylogenetic trees, representing 84% of the isolates from these lineages. A similar analysis restricted to pairs of isolates from the same herd showed a similar peak of closely related isolates with 98 isolates with less than five differences with another isolate out of the 108 isolates with less than 21 different alleles (Fig. 4B, Table S4). To estimate the rate of dissemination we computed the genetic distance between isolates as the number of different alleles by cgMLST and the geographic distance between farms (Fig. 4c). We observed two jumps in the distance distribution. Isolates from the 254 pairs differing by less than five alleles were at a median distance of 10 km and 203 (80%) were from herds less than 25 km apart with. This suggested local spread of resistant *E. coli* between these farms. Isolates from pairs differing by 5 to 20 alleles were at a median distance of 49 km compared to 66 km for isolates differing by over 20 alleles. This observation suggests other longer-term constraints on *E. coli* dissemination.

**Fig. 4.**
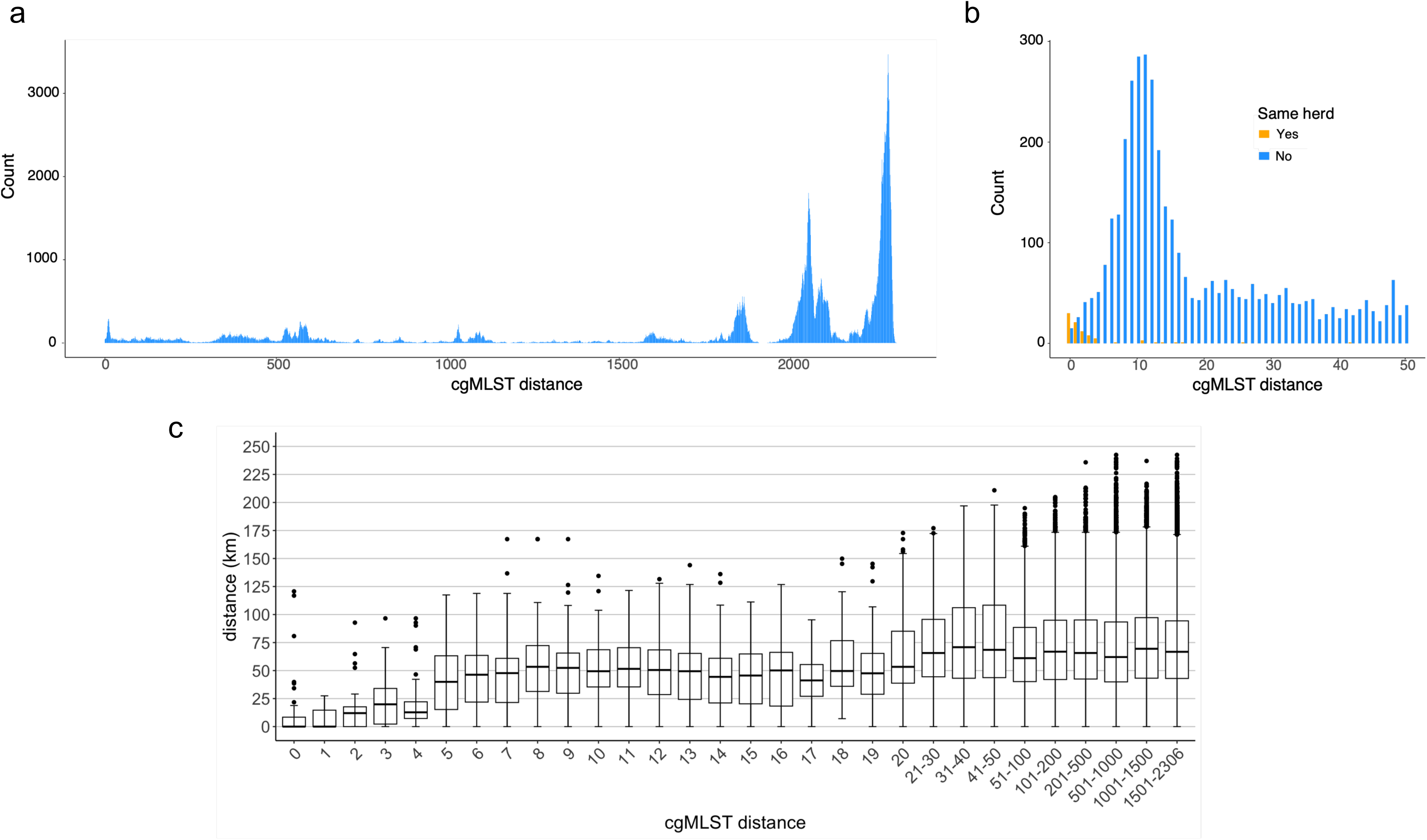
Genetic and geographic distances between isolates. **a.** distribution of pairwise genetic distances in number of different cgMLST alleles for all pairs of deduplicated isolates from different herds. **b.** Same as A, but for pairs with 0 to 50 different alleles. In orange, pairs of isolates from the same herd. **c**. Relation between the genetic distance in number of different alleles and the physical distance in km.

To further analyze isolate dissemination, we geographically localized those belonging to 11 lineages with more than nine isolates from our collection. Isolates closely related in some lineages clustered geographically. This is the case for sub-clusters of isolates in L-ST-88-1 (n=6), L-ST88-2 (n=7) and L-ST88-3 (n=3) (Fig. 5), and in four other STs: 167, 617, 1642 (Fig. S9) and ST10 (Fig. S10). Our definition of lineage was independent of the evolutionary branch length, explaining why local dissemination was variable among these lineages. This analysis revealed, in addition to the geographic proximity of isolates showing low genetic distance, differences between the east and west of the river Meuse. All isolates from lineages L-ST88-1 and L-ST88-3 were from herds located in the eastern part of Wallonia, two ST783 sublineages were restricted to the west of the Meuse, whereas the third one was mainly in the east (Fig. 5d). This geographic distribution might correspond to the jump in distances observed for pairs with over four different alleles (Fig. 4c)

**Fig. 5.**
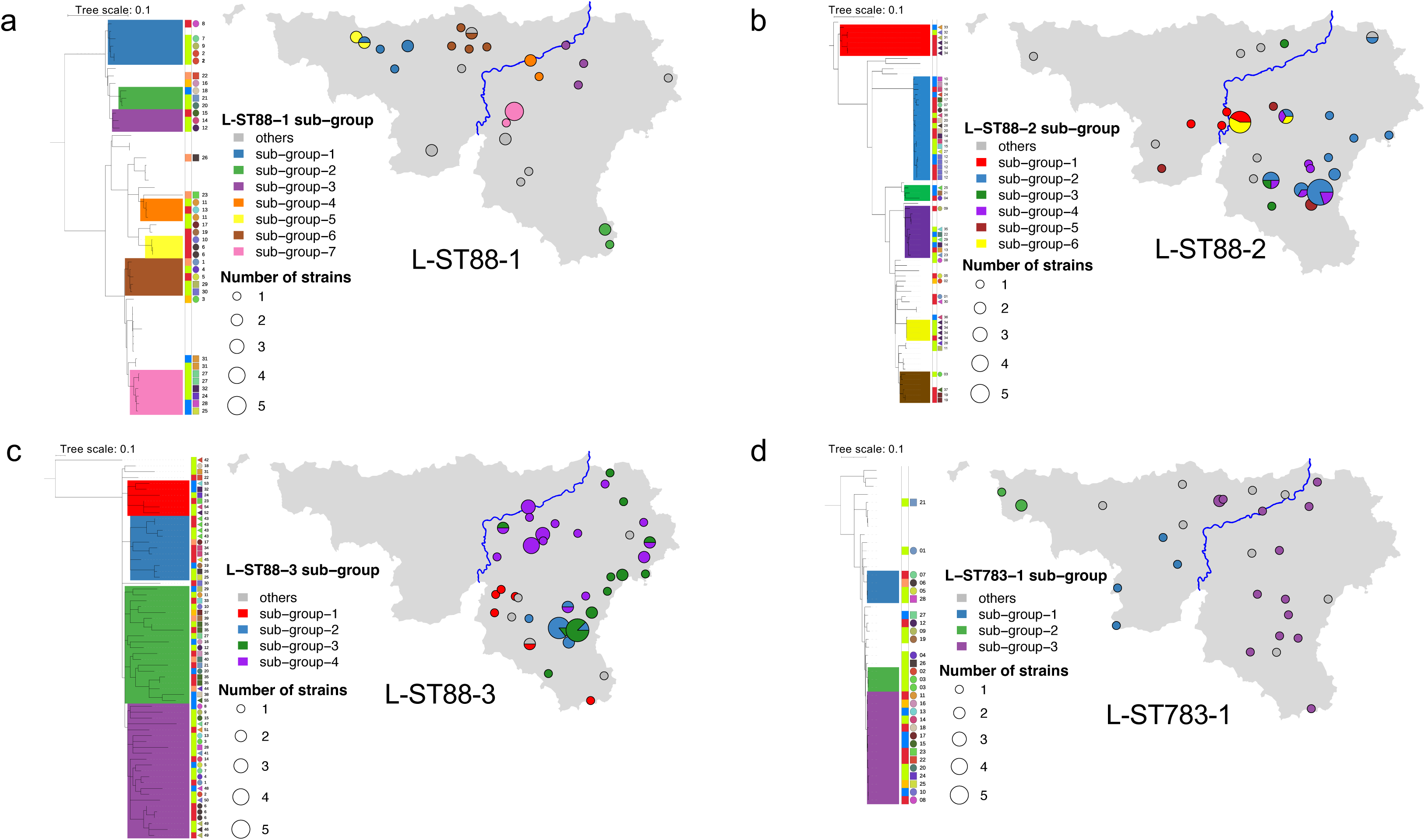
Local dissemination of closely related isolates and geographic specificities. The geographic localization was compared to the phylogenetic proximity by localizing isolates according to the GPS coordinates of the postal code. Size of the circle corresponds to the number of isolates. Colors correspond to tree sublineages and as sectors if isolates from several sub-lineage. Isolates were annotated on the right of the tree according to the figure key. From left to right, the season (in white isolates from EnteroBase) and the herd. The blue line represents the Meuse. **a.** L-ST88-1, isolates were of the whole Wallonia with local clustering of seven sub-lineages. **b.** Lineages L-ST88-2, 34 isolates where localized east of the Meuse and seven west. **c.** Lineage L-ST88-3 all strains were isolated east of the Meuse. **d.** Lineage L-ST783-1 isolates were of the whole Wallonia with local clustering of three sub-lineages

To further quantify the impact of the geography on transmission, we analyzed the geographic distribution of isolates from seasons S1-3 (Fig. 6a). We received and phenotyped 813 isolates from 445 herds from the more agricultural eastern region and 654 isolates from 370 herds in the west (Tables S1, S5). 26 isolates were from 14 farms located in postal codes crossed by the Meuse. The distribution of β-lactam resistance classes was significantly different between east and west of the Meuse. The frequency of AmpC-like isolates was twice as high in the east as in the west, while the ESBL isolates were more frequent in the west. The proportion of NSBL and AmpC classes were similar (Table S5). These differences were also observed for sequenced isolates from the five calving seasons (Table S5). In agreement with the dominance of AmpC-like profiles in the east, we observed a more than double proportion of ST88 isolates. However, the proportion of ST88 isolates not part of a cluster was similar between east and west, with 2.7% and 2.8% of the sequenced isolates respectively (Table S5).

**Fig. 6:**
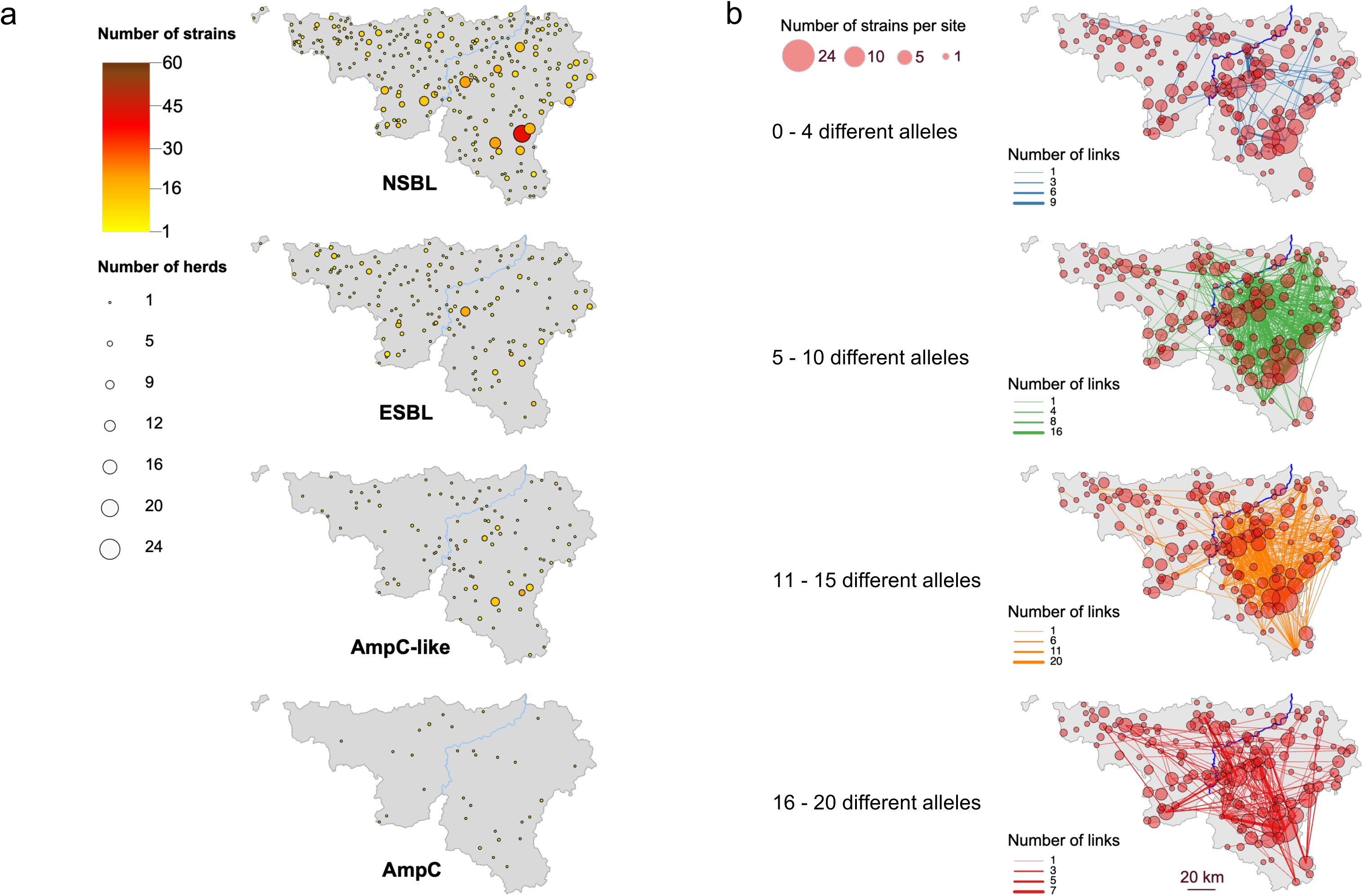
ß-lactam resistance profiles and transmission differences between west and east of the Meuse. **A**. Geographic distribution of the isolates belonging to the four classes of ß-lactam resistance. In each postal code, the number of isolates and the number of herds were indicated by the color and the circle diameter respectively according to the figure key. **B**. Geographic link of isolates pairs showing less than five different alleles by cgMLST (blue links), between 5 and 10 different alleles (green links), between 11 and 15 (orange links) and between 16 and 20 (red links). The circle diameter represents the number of isolates per postal code and the line thickness the number of isolates pairs according to the figure key.

Next, we analyzed the geographic distribution of isolate pairs between farms from the five calving seasons showing less than 21 different alleles by cgMLST (Fig. 6b, Table S4). We considered all isolates, only ST88 isolates, only isolates with a AmpC-like profile and conversely all isolates except ST88 or except AmpC-like profiles and performed the statistical analysis on the number of isolates belonging to at least one pair (Table S6). After normalization by the number of isolates, we observed ten times more pairs with less than 21 different alleles on the east compared to the west of the Meuse. This was largely attributable to isolates with an AmpC-like profile or from ST88. However, the difference between east and west was still significant when excluding AmpC-like or ST88 isolates. East-west isolate pairs were mainly non-AmpC like or non-ST88 and were more frequent than west-west pairs. When considering transmissions within a farm, there were no difference between east and west. These findings confirmed the observation made on individual clusters. The local geography (east or west of the Meuse) had a strong impact on the phenotype of the collected isolates and on *E. coli* dissemination.

## Discussion

*E. coli* is a major cause of deadly infections in calves, resulting in economic losses for farmers. Thanks to our extensive sampling of *E. coli* in sick calves throughout Wallonia, we were able to answer key questions that are important for better managing *E. coli* infections in calf rearing and assess their zoonotic potential. Isolates were collected from 444 farms scattered across Wallonia and isolated over five calving seasons from 2014 to 2020.

The restriction of antibiotics essential for human health in Belgium^11^ led to a reduction in 3GC-and 4GC-resistant isolates (described as ESBL and AmpC phenotypes), compensated by an increase in isolates still resistant to amoxicillin but not to 3GC or 4GC^12^. Despite the impact on our selection for WGS of a balanced but non-proportional collection, we showed that, considering the four β-lactam resistance profiles during the three calving seasons from 2017 to 2020, there was no significant change in the population structure based on ST, nor a significant impact on β-lactamase gene frequencies (Fig. 2). This stability suggests that antibiotics select for resistance in a bacterial population adapted to a host and do not shape the population.

One specificity of our collection was the origin of the isolates, from sick or dead calves, which showed resistance to ampicillin. Although the isolates were selected on this single resistance criteria, which is common among *E. coli*, all except nine were predicted to be MDR. Even isolates with the NSBL profile carried a median of nine ARGs. Phylogenetic analyses showed that most isolates clustered at different phylogenetic levels. They belong mainly to STs reported to be associated with bovine hosts, notably calves, like ST88, ST10, ST117, ST167, ST69 and ST783 in decreasing order of incidence^23–25^ (Fig. 1 and 2). Within these STs, 54% of the isolates clustered in sublineages with more than five isolates and a minority of isolates from EnteroBase. EnteroBase isolates were also MDR (Table S3). These groupings probably reflect a combination of bovine (calf) adaptation and local dissemination. 93% of EnteroBase isolates belonging to these clusters were from European countries. This suggests a restricted reservoir of circulating MDR isolates possibly contributing to the observed stability of the population.

Half of the EnteroBase isolates from these clusters were of human origin. Two human isolates carrying the *bla*_OXA-48_ carbapenemase gene belonged to clusters L-ST88-2 and L-ST88-5, respectively. As we predicted mainly transitions from bovine to human hosts, these highly resistant lineages exhibit zoonotic potential, and cattle herds might represent a long-term reservoir for human MDR *E. coli* strains.

ST88 represented almost 30% of all sequenced isolates in our study. ST88 isolates have been frequently isolated from bovine hosts and were previously reported as mutated in P*_ampC_* both in animals and humans^24,26^, however, not on such a scale. Our sampling of isolates with an AmpC-like profile representing 25% of the sequenced isolates and 61% of ST88 isolates contributed to this high proportion. We also observed a strong association between derepression of *ampC* and ST88 with 75% of the 223 strains mutated in P*_ampC_* belonging to this ST and 75% of the ST88 isolates were mutated in P*_ampC_*. This association could result from a higher level of resistance due to AmpC overproduction (better selection) or from a lower fitness cost (better dissemination) in ST88 strains. The AmpC-like phenotype resulting from mutations in P*_ampC_* implies a reduced susceptibility to the amoxicillin/clavulanate combination. The restriction on the use of critical antibiotics, including 3GC and 4GC, resulted in an increased use of this combination, which has been suggested to contribute to an increased incidence of AmpC-hyperproducing *E. coli*^27^. Furthermore, the high proportion of ST88 isolates in our study resulted mainly from the dissemination of a few lineages mutated in P_ampC_ in the eastern part of Wallonia. Therefore, the high proportion of ST88/ST783 might result from (i) selection of P*_ampC_* mutations due to the use of amoxycillin/clavulanate in livestock, (ii) adaptation of ST88/ST783 to the bovine host, and (iii) the stability of a phenotype due to a chromosomal mutation. ST88 isolates are recognized extraintestinal pathogenic *E. coli*, which have been associated with a high mortality rate in human bloodstream infections^28^. ST88 isolates from sick animals might similarly show a high capacity to cause disease in calves. However, to our knowledge there is no report strengthening this hypothesis.

ST10 was the second most abundant ST with 81 isolates over the five calving seasons. ST10 is the most frequent ST reported in EnteroBase. Surprisingly, 89% of the ST10 isolates from the study clustered with a cgMLST distance lower than 100 together with only 2.6% of ST10 isolates from EnteroBase. Enterobase isolates with reported origin were mainly from animals (182 versus 63 from humans) (Fig. S3a). Compared to the bulk of ST10 strains, this cluster might be particularly adapted to the bovine host. The selection of isolates from sick animals might also have contributed to the overrepresentation of this lineage. The high frequency of ST117, the third most abundant lineage with 72 isolates during the five seasons was unexpected, since ST117 is an avian pathogenic lineage associated with poultry^29,30^. Of the 1525 genome sequences retrieved from EnteroBase, only 56 were from bovine isolates compared to 1122 from bird or poultry and 122 from humans. 89% of the ST117 isolates from our study belong to three sublineages together with seven isolates from EnteroBase: four from humans, one from bovine and one from swine, but none from birds. These three lineages might have acquired an adaptation to the bovine host and subsequently be transmitted to other hosts. ST117 is frequently pathogenic in birds and in humans^29,30^. The sampling of isolates from sick animals might have contributed to its high frequency.

The transmission of bacteria between farms can be caused either by the transfer of animals, or by indirect contacts with the environment via vehicles traveling between farms or other fomites. For short evolutionary distances (a difference of less than five alleles), isolates were clustered geographically with a median farm distance of 10 km (Fig. 4). The geographic localization of phylogenetic clusters confirmed the local transmission of closely related isolates (Fig. 5, Fig. S9, S10). Animal transfers between farms are expected to be rare as calf rearing farms are mainly (94%) localized in the Flemish region of Belgium^31^. The local transmission via fomite by indirect contact is therefore likely the main route of transmission. In the context of dairy farms, *E. coli* clones are transmitted first within a farm and then very locally, before further spreading. Dissemination of Shiga toxin-producing *E. coli* O157:H7 among dairy farms in close geographic proximity was similarly described in Northeast Ohio, USA^32^.

We observed very different behavior between two areas in Wallonia separated by the Meuse. Cluster of geographically and genetically closely related isolates were abundant in the east. While transmissions were much less frequent in the west and between east and west (Fig. 6). This difference was mainly due to the geographic distribution of AmpC-like clusters dominant in the east and their propension to disseminate. However, even when AmpC-like or ST88 isolates were not considered, pairs of isolates on east were more frequent than on west (Table S6). Conversely, ESBL isolates were more abundant in the west. Different factors might contribute to the observed differences between the two parts of Wallonia, which might include different antibiotic usage and farming practices^33,34^.

We acknowledge some limitations in our study. Only β-lactam resistant isolates were analyzed. However, the rate of ampicillin-susceptible isolates was low, ranging from 11.1 to 16%^12^ during the study period. Therefore, our analysis remains largely representative of the circulating strains and provide an accurate view on resistant isolates. In this study, we have only analyzed isolates from sick animals. A similar analysis of carriage isolates from healthy animals would allow us to identify specificities of the population in sick animals. We also did not analyze *E. coli* isolated from the environment to confirm its contribution to the between farms dissemination of MDR *E. coli*. Our systematic sampling started in November 2017 and the restriction on 3GC started in 2016. However, reduction of 3GC resistance was also observed during the three seasons we have analyzed^12^. Antibiotic usage per herd or per postal code would allow to determine its contribution to the difference between west and east of the Meuse. However, these data are not available to our knowledge.

Our analysis provides clues on the resilience of calf-infecting *E. coli* in terms of population and β-lactam resistance mechanisms. It also suggests the transmission from bovines to humans, and the zoonotic potential of MDR lineages. The extensive geographic sources of the isolates demonstrate the importance of local transmission of *E. coli* and reveals striking differences between the east and west of Wallonia. It provides means to reduce transmission by limiting the transfer of contaminated material between closely related farms as done prior to limit transmission of the viral foot and mouth disease^35^.

## Material and methods

### *E. coli* isolates, growth conditions and antibiotic susceptibility testing

The 1495 *E. coli* from seasons S1-3 were isolated from the feces of diarrheic calves (n=962) and from internal organs (n=272) or the intestinal content (n=261) of septicemic calves necropsied at ARSIA (Table S1). Isolates from internal organs or with positive agglutination test for fimbriae F5 or F17, the CS31 antigen (Biovac, France), or enterohemolysin-producers on washed sheep blood agar plates were considered pathogenic^36^. The 362 remaining isolates from feces or intestinal content were selected on McConkey agar plates complemented with 1 mg/L cefotaxime (Led Techno, The Netherlands). Antibiotic susceptibility testing for resistance against eight β-lactams was performed by disk (I2A, France) diffusion assay following the EUCAST/CASFM-vet guidelines (Table S1). Isolates were classified into four β-lactam resistance profiles: “Narrow-Spectrum-β-lactamase” (NSBL) amoxycillin resistant only, “Extended-Spectrum-β-lactamase” (ESBL) 3GC resistant with clavulanic acid synergy, “Cephalosporinase” (AmpC) 3GC resistant without clavulanic acid synergy, and AmpC-like amoxycillin resistant with no or reduced clavulanic synergy and cefoxitin resistant^12,37^ (Table 1, Tables S1).

681 isolates were selected for WGS to keep the balance between the four β-lactam resistance profiles across the three calving seasons. They were from 379 farms spread out over Wallonia (Table 1, Tables S1). There were up to 13 farms per postal code and 1 to 11 sequenced isolates per farm (Fig. S1). Two isolates from different anatomic origins were selected from 68 calves and three isolates from one calf. To obtain more temporal data, we selected 83 additional isolates, considered pathogenic according to our criteria from calving seasons 2014-2015 and 2015-2016 (aS1, aS2) (Table S2). Fifteen were from herds already sampled during seasons S1-3 and 68 from 65 additional farms making a total of 444 herds. To respect laws on data privacy, animal and herd identifications were encoded at the data source before analysis.

### Whole Genome Sequencing, sequence assembly and analyses

DNA extractions were performed with the Qiagen DNeasy Blood & Tissue Kit. Sequencing libraries were prepared following the manufacturer’s protocol (NEBNext Ultra II FS DNA library prep kit). WGS was performed using a NextSeq500 sequencer (Illumina) in pair-ended reads (2 x 75 nucleotides) or a Novaseq (2 x 150 nucleotides). Reads were processed with Cutadapt 2.10^38^ and Trimmomatic 0.39^39^. *De novo* assembly was performed using Shovill 1.1.0 (RRID:*SCR_017077*). All sequences are available under the NCBI bioproject PRJEB79574^37^ and PRJNA566319 (Tables S1, S2).

Multilocus Sequence Typing (MLST), based on the Achtman scheme, was done with mlst 2.19.0 (https://github.com/tseemann/mlst), and Clermont Typing was performed with the *in silico* method^40^. Resfinder 4^41^ and Pointfinder^42^ were performed via ABRicate 1.0.1 to detect resistance genes and mutations^43^. Draft genomes were annotated using Prokka 1.14.5^44^. Core-genome MLST (cgMLST) was performed with chewBBACA 3.3.6^45^, based on the RIDOM *E. coli* scheme. Numbers of alleles different between two isolates were determined with cgMLST-dists 0.4.0 (https://github.com/tseemann/cgmlst-dists).

### Phylogenetic analyses

A maximum likelihood (ML) core genome phylogeny of the 764 isolates was determined with Roary, 3.13.0^46^ and RAxML 8.2.12^47^, with *Escherichia fergusonii* (NZ_CP057657) as outgroup. Genomes from the main sequence types (STs) were contextualized with EnteroBase datasets downloaded in July 2022^18^. Only the 1521 ST117 sequences with complete metadata were retained. A preliminary analysis showed that 72 out of the 81 ST10 isolates belong to the cgMLST hierarchical cluster HC100 1126. We included only the 353 genomes from this HC100 out of the 13592 ST10 Enterobase isolates. ML phylogenies were generated from recombination-filtered core alignments with RAxML^47^, using Snippy v4.6.0 (https://github.com/tseemann/snippy) and Gubbins v2.4.1^48^ and visualized with iTOL v7^49^. The ancestral state (bovine or human) at each node of phylogenies was inferred using PastML v1.9.51 using the MPPA (Maximum Parsimony with Posterior Averaging) prediction method and the F81 model^22^. ML phylogenies for the PastML analysis were inferred with IQ-TREE/2.4.0, using the previous alignments and the GTR+F+ASC+R2 model^22^. Branch support was assessed via 1000 ultrafast bootstrap replicates, and topologies refined by BNNI^50^.

### Geographical distribution of the isolates and clonal dissemination

Isolates and farms were localized according to the GPS coordinates of their postal codes (Open Data Wallonie-Bruxelles, https://www.odwb.be/). Farms were classified as east, west, or crossed by the Meuse. After deduplicating clonal isolates from the same animal, we calculated genetic (cgMLST allelic) and geographic distances between pairs. Maps and network visualizations were generated in R using standard spatial packages (R Core Team, https://www.r-project.org/).

### Statistical analysis

Statistical analyses of isolate characteristics were performed with R and Python3 using Fisher’s exact test or the chi-squared test, as appropriate. The difference in the ARG number between *mcr*-carrying and non-*mcr*-carrying isolates was assessed using the non-parametric Wilcoxon rank-sum test for independent samples.

## Data availability

All sequences are available under the NCBI bioproject PRJEB79574^37^ and PRJNA566319 (Tables S1, S2). Isolate and sequence data are in Table S1 to S3. Geographic and genetic distances between isolate pairs are in Table S4.

## Acknowledgement

We thank Patrick Mayrens, Edouard Reding and Bernard Christiaens for information on calf rearing in Belgium and Marisa Haenni and Jean-Yves Madec for information on *E. coli* in livestock. Sequencing was performed at the Biomics Platform, C2RT, Institut Pasteur, Paris, France, supported by France Génomique (ANR-10-INBS-09) and IBISA. This work was supported by the “Investissement d’Avenir” programs LABEX IBEID (Grant ANR-10-LABX-62-IBEID), DYASPEO (ANR-20-PAMR-003) and SEQ2DIAG (ANR-20-PAMR-0010). Virginie Guérin’s internship at the Institut Pasteur was supported by the Belgian Federal Public Service Health, Food Chain Safety and Environment, Grant n° RF 17/6317 RU-BLA-ESBL-CPE and by the Fédération Wallonie-Bruxelles, Grant n° CHK/MA/JCD/CBV 20-41.

## Author contributions

V.G., J.G.M and P.G. conceived the study; V.G., J.-N.D and N.C. performed the experiments;

N.C. and V.G. carried out bioinformatics analyses, P.G., V.G., G.R. and N.C. analyzed the data;

D.T. and M.S. provided materials. V.G., P.G., G.M.M.D.M. and N.C. wrote the manuscript. All authors corrected the manuscript.

## References

1 Denamur, E., Clermont, O., Bonacorsi, S. & Gordon, D. The population genetics of pathogenic *Escherichia coli*. Nat Rev Microbiol 19, 37–54 (2021). 10.1038/s41579-020-0416-x

2 Ramos, S., et al. *Escherichia coli* as Commensal and Pathogenic Bacteria Among Food-Producing Animals: Health Implications of Extended Spectrum β-lactamase (ESBL) Production. Animals (Basel) 10 (2020). 10.3390/ani10122239

3 Kolenda, R., Burdukiewicz, M. & Schierack, P. A systematic review and meta-analysis of the epidemiology of pathogenic *Escherichia coli* of calves and the role of calves as reservoirs for human pathogenic *E. coli*. Frontiers in cellular and infection microbiology 5, 23 (2015). 10.3389/fcimb.2015.00023

4 Cho, Y. I. & Yoon, K. J. An overview of calf diarrhea - infectious etiology, diagnosis, and intervention. J Vet Sci 15, 1–17 (2014). 10.4142/jvs.2014.15.1.1

5 Uyama, T. et al. Associations of calf management practices with antimicrobial use in Canadian dairy calves. J Dairy Sci 105, 9084–9097 (2022). 10.3168/jds.2022-22299

6 Holmes, A. H. et al. Understanding the mechanisms and drivers of antimicrobial resistance. Lancet 387, 176–187 (2016). 10.1016/s0140-6736(15)00473-0

7 Velazquez-Meza, M. E., Galarde-López, M., Carrillo-Quiróz, B. & Alpuche-Aranda, C. M. Antimicrobial resistance: One Health approach. Vet World 15, 743–749 (2022). 10.14202/vetworld.2022.743-749

8 Jesudason, T. WHO publishes list of medically important antimicrobials. Lancet Infect Dis 24, e284 (2024). 10.1016/s1473-3099(24)00248-2

9 Aidara-Kane, A. et al. World Health Organization (WHO) guidelines on use of medically important antimicrobials in food-producing animals. Antimicrobial resistance and infection control 7, 7 (2018). 10.1186/s13756-017-0294-9

10 Tiseo, K., Huber, L., Gilbert, M., Robinson, T. P. & Van Boeckel, T. P. Global Trends in Antimicrobial Use in Food Animals from 2017 to 2030. *Antibiotics (Basel*, Switzerland) 9 (2020). 10.3390/antibiotics9120918

11 Royal Decree of Belgium. Arrêté Royal du 21 Juillet 2016. Arrêté Royal Relatif aux Conditions d’Utilisation des Médicaments par les Médecins Vétérinaires et par les Responsables des Animaux. Available online: http://www.etaamb.be/fr/arrete-royal-du-21-juillet-2016_n2016024152.html. (In French).

12 Guérin, V. et al. Seven-Year Evolution of β-Lactam Resistance Phenotypes in *Escherichia coli* Isolated from Young Diarrheic and Septicaemic Calves in Belgium. Veterinary sciences 9 (2022). 10.3390/vetsci9020045

13 Andersson, D. I. & Hughes, D. Selection and Transmission of Antibiotic-Resistant Bacteria. Microbiology spectrum 5 (2017). 10.1128/microbiolspec.MTBP-0013-2016

14 Croucher, N. J. & Didelot, X. The application of genomics to tracing bacterial pathogen transmission. Curr Opin Microbiol 23, 62–67 (2015). 10.1016/j.mib.2014.11.004

15 Ludden, C. et al. One Health Genomic Surveillance of *Escherichia coli* Demonstrates Distinct Lineages and Mobile Genetic Elements in Isolates from Humans versus Livestock. mBio 10 (2019). 10.1128/mBio.02693-18

16 De Koster, S. et al. One Health surveillance of colistin-resistant Enterobacterales in Belgium and the Netherlands between 2017 and 2019. PLoS One 19, e0298096 (2024). 10.1371/journal.pone.0298096

17 Milenkov, M. et al. Implementation of the WHO Tricycle protocol for surveillance of extended-spectrum β-lactamase producing *Escherichia coli* in humans, chickens, and the environment in Madagascar: a prospective genomic epidemiology study. Lancet Microbe 5, 100850 (2024). 10.1016/s2666-5247(24)00065-x

18 Zhou, Z., Alikhan, N. F., Mohamed, K., Fan, Y. & Achtman, M. The EnteroBase user’s guide, with case studies on *Salmonella* transmissions, *Yersinia pestis* phylogeny, and *Escherichia* core genomic diversity. Genome Res 30, 138–152 (2020). 10.1101/gr.251678.119

19 Clermont, O. et al. Characterization and rapid identification of phylogroup G in *Escherichia coli*, a lineage with high virulence and antibiotic resistance potential. Environmental microbiology 21, 3107–3117 (2019). 10.1111/1462-2920.14713

20 Caroff, N., Espaze, E., Gautreau, D., Richet, H. & Reynaud, A. Analysis of the effects of −42 and −32 *ampC* promoter mutations in clinical isolates of *Escherichia coli* hyperproducing *ampC*. J Antimicrob Chemother 45, 783–788 (2000). 10.1093/jac/45.6.783

21 Mulvey, M. R. et al. Molecular characterization of cefoxitin-resistant *Escherichia coli* from Canadian hospitals. Antimicrob. Agents Chemother. 49, 358–365 (2005).

22 Ishikawa, S. A., Zhukova, A., Iwasaki, W. & Gascuel, O. A Fast Likelihood Method to Reconstruct and Visualize Ancestral Scenarios. Mol Biol Evol 36, 2069–2085 (2019). 10.1093/molbev/msz131

23 de Lagarde, M. et al. Clonal and plasmidic dissemination of critical antimicrobial resistance genes through clinically relevant ExPEC and APEC-like lineages (ST) in the dairy cattle population of Québec, Canada. Frontiers in microbiology 14, 1304678 (2023). 10.3389/fmicb.2023.1304678

24 Haenni, M., Châtre, P. & Madec, J. Y. Emergence of *Escherichia coli* producing extended-spectrum AmpC β-lactamases (ESAC) in animals. Frontiers in microbiology 5, 53 (2014). 10.3389/fmicb.2014.00053

25 Masarikova, M. et al. Genomic analysis of extended-spectrum beta-lactamase-producing *E. coli* from Czech diary calves and their caretakers. Front Vet Sci 12, 1552297 (2025). 10.3389/fvets.2025.1552297

26 Alzayn, M. et al. Characterization of AmpC-hyperproducing *Escherichia coli* from humans and dairy farms collected in parallel in the same geographical region. J Antimicrob Chemother 75, 2471–2479 (2020). 10.1093/jac/dkaa207

27 Burgess, S. A. et al. The epidemiology of AmpC-producing *Escherichia coli* isolated from dairy cattle faeces on pasture-fed farms. J. Med. Microbiol. 70 (2021). 10.1099/jmm.0.001447

28 de Lastours, V. et al. Mortality in *Escherichia coli* bloodstream infections: antibiotic resistance still does not make it. J Antimicrob Chemother 75, 2334–2343 (2020). 10.1093/jac/dkaa161

29 Xia, F. et al. Complete genomic analysis of ST117 lineage extraintestinal pathogenic *Escherichia coli* (ExPEC) to reveal multiple genetic determinants to drive its global transmission: ST117 *E. coli* as an emerging multidrug-resistant foodborne ExPEC with zoonotic potential. Transbound Emerg Dis (2022). 10.1111/tbed.14678

30 Ronco, T. et al. Spread of avian pathogenic *Escherichia coli* ST117 O78:H4 in Nordic broiler production. BMC Genomics 18, 13 (2017). 10.1186/s12864-016-3415-6

31 Celagri. [Key figures] The dairy calf fattening sector (In French, [Chiffres clés] La filière d’engraissement des veaux laitiers) https://www.celagri.be/wp-content/uploads/2024/02/La-filiere-veau-chiffres-cles-2022.pdf. (2024).

32 Wetzel, A. N. & LeJeune, J. T. Clonal dissemination of *Escherichia coli* O157:H7 subtypes among dairy farms in northeast Ohio. Appl. Environ. Microbiol. 72, 2621–2626 (2006). 10.1128/aem.72.4.2621-2626.2006

33. Cheptel bovin laitier - Etat de l’Agriculture Wallonne (Dairy cattle herd - State of Walloon Agriculture), Available online file:///Users/pglaser/Downloads/Bovins_laitiers.pdf (In French). (2024).

34. Cheptel bovin viandeux - Etat de l’Agriculture Wallonne (Beef cattle herd - State of Walloon Agriculture) file:///Users/pglaser/Downloads/Bovins_viandeux.pdf (in French). (2024).

35 Ellis, J., Brown, E., Colenutt, C., Schley, D. & Gubbins, S. Inferring transmission routes for foot-and-mouth disease virus within a cattle herd using approximate Bayesian computation. Epidemics 46, 100740 (2024). 10.1016/j.epidem.2024.100740

36 Guérin, V. et al. A three-year evolution and comparison of the bla(CTX-M) genes in pathogenic and non-pathogenic *Escherichia coli* isolated from young diarrheic and septicaemic calves in Belgium. Research in veterinary science 152, 647–650 (2022). 10.1016/j.rvsc.2022.09.037

37 Guérin, V. et al. Identification of β-Lactamase-Encoding (bla) Genes in Phenotypically β-Lactam-Resistant *Escherichia coli* Isolated from Young Calves in Belgium. Microb. Drug Resist. 27, 1578–1584 (2021). 10.1089/mdr.2020.0472

38 Martin, M. Cutadapt removes adapter sequences from high-throughput sequencing reads. EMBnet J., 10 (2011).

39 Bolger, A. M., Lohse, M. & Usadel, B. Trimmomatic: a flexible trimmer for Illumina sequence data. Bioinformatics 30, 2114–2120 (2014). 10.1093/bioinformatics/btu170

40 Beghain, J., Bridier-Nahmias, A., Le Nagard, H., Denamur, E. & Clermont, O. ClermonTyping: an easy-to-use and accurate in silico method for *Escherichia* genus strain phylotyping. Microbial genomics 4 (2018). 10.1099/mgen.0.000192

41 Bortolaia, V. et al. ResFinder 4.0 for predictions of phenotypes from genotypes. J Antimicrob Chemother (2020). 10.1093/jac/dkaa345

42 Zankari, E. et al. PointFinder: a novel web tool for WGS-based detection of antimicrobial resistance associated with chromosomal point mutations in bacterial pathogens. J Antimicrob Chemother 72, 2764–2768 (2017). 10.1093/jac/dkx217

43 Zankari, E. et al. Identification of acquired antimicrobial resistance genes. J Antimicrob Chemother 67, 2640–2644 (2012). 10.1093/jac/dks261

44 Seemann, T. Prokka: rapid prokaryotic genome annotation. Bioinformatics 30, 2068–2069 (2014). 10.1093/bioinformatics/btu153

45 Silva, M. et al. chewBBACA: A complete suite for gene-by-gene schema creation and strain identification. Microbial genomics 4 (2018). 10.1099/mgen.0.000166

46 Page, A. J. et al. Roary: rapid large-scale prokaryote pan genome analysis. Bioinformatics 31, 3691–3693 (2015). 10.1093/bioinformatics/btv421

47 Stamatakis, A. RAxML version 8: a tool for phylogenetic analysis and post-analysis of large phylogenies. Bioinformatics 30, 1312–1313 (2014). 10.1093/bioinformatics/btu033

48 Croucher, N. J. et al. Rapid phylogenetic analysis of large samples of recombinant bacterial whole genome sequences using Gubbins. Nucleic Acids Res. 43, e15 (2015). 10.1093/nar/gku1196

49 Letunic, I. & Bork, P. Interactive Tree Of Life (iTOL) v5: an online tool for phylogenetic tree display and annotation. Nucleic Acids Res. 49, W293–w296 (2021). 10.1093/nar/gkab301

50 Minh, B. Q. et al. IQ-TREE 2: New Models and Efficient Methods for Phylogenetic Inference in the Genomic Era. Mol Biol Evol 37, 1530–1534 (2020). 10.1093/molbev/msaa015

