## Supplementary tables and figures for "Spatiotemporal dynamics of β-lactam-resistant *E. coli* in young diseased calves in Wallonia, Belgium"

**Saptio-temporal dynamic of β-lactam resistant *E. coli* in young diseased calves in Wallonia, Belgium.**

### Supplementary material.

**Table S1: Characteristics of *E. coli* isolates collected during the three calving seasons, S1, S2 and S3.** Excel file Table S1

[§]age: 0 means the first day of life; [&]side: East or West of the Meuse, Meuse indicates that the Meuse crosses the postal code; [#]Other isolate from the same animal: no, means that there is no other isolate from the same animal in our collection, a star indicates that the isolates from this animal are clonal (less than five different alleles by cgMLST), [$]P/NP: P is for pathogenic and NP for non-pathogenic as defined in the main text. [€]Minca: the positive antigen is indicated; N for negative. [ß]Ehly negative (N) or positive (P) for Enterohemolysin activity, H+ is for alpha-hemolysin, ? is for doubtful result; *McConkey-CTX: positive (P) or negative (N) for growth on Mac Conkey plate supplemented with 1 mg/L cefotaxime, ? is for doubtful result; [%]Lineage: details on lineages are in Table 2. [£]Antibiotic susceptibility testing, inhibition zone diameter and Susceptible (S), Intermediate (I), or Resistant (R) phenotype according to EUCAST breakpoints. ND, not determined. AMX, Amoxicillin, AMC, Amoxicillin / Clavulanate; XNL, Ceftiofur; CFQ cefquinome, CTX cefotaxime; CTC, cefotaxime / Clavulanate; FOX, cefoxitin; MER, meropenem.

**Table S2: Characteristics of *E. coli* isolates collected during the two calving seasons, aS1 and aS2** Excel file Table S2

[&]side: East or West of the Meuse. Meuse indicates that the Meuse crosses the postal code; [#]Other isolate from the same animal: no, means that there is no other isolate from the same animal in our collection. [%]Lineage: details on lineages are in Table 2.

**Table S3**: **Characteristics of *E. coli* EnteroBase isolates clustering with isolates from the study**. Excel file Table S3

[§]ID Enterobase identifiers. Annotations were retrieved from Enterobase

**Table S4: Genetic and geographic distances of isolates pairs with less than 21 different cgMLST alleles.** Excel file Table S4

[&]Genetic distances in number of different cgMLST alleles. #Meuse means that the postal code is crossed by the river Meuse.

**Table S5** Geographical and phenotypic distribution of *E. coli* isolates

| | Strain collection (S1-3) | | | | | Deduplicated sequenced isolates (aS1-2, S1-3) | | | | |
|---|---|---|---|---|---|---|---|---|---|---|
| | All strains | All NSBL | All ESBL | AmpC-like | AmpC | All strains | AmpC-like | ST88 | ST88-Cluster (%ST88, % all) | ST88 other |
| East | 813 | 449 (55,2%) | 184 (22,6%) | 147 (18%) | 29 (3,6%) | 415 | 128 | 148 (18,1%) | 126 (85%, 15,4%) | 22 (2,7%) |
| West | 654 | 375 (57,3%) | 195 (29,8%) | 59 (9%) | 21 (3,2%) | 306 | 51 | 55 (8,4%) | 37 (67,3%, 5,7%) | 18 2,8% |
| [§]Meuse | 26 | 13 (50%) | 8 (30,8%) | 5 (19,2%) | 0 | 15 | 5 | 4 (15,4%) | 2 (50%, 7.7%) | 2 (7.7%) |
| P value[&] | | 0.43 | [@]0.0022 | <0.0001 | 0.67 | | <0.0001 | <0.0001 | 0.0089 (ST88) <0.0001 (all) | [*]0.089 (ST88) 0,74 (all) |

[§]Herds in postal codes crossed by the Meuse. [&]Two-tailed exact Fisher test, East compared to West. [@]P value that ESBL are more frequent on the West compared to the East. [*]P value that ST88 are more frequent outside cluster on the west compared to the east.

**Table S6** Isolate pairs with less than 21 different alleles and isolates with at least one match

| | [#]All strains | [§]P values | [#@]AmpC-Like | [§]P values | [#]No AmpC-Like | [§]P values | [#]ST88 | [§]P value | [#]No ST88 | [§]P valuer |
|---|---|---|---|---|---|---|---|---|---|---|
| [€]East <=> East | 2130 (281) | - | 1875 (110) | - | 255 (171) | - | 1907 (124) | | 223 (157) | |
| [€]West <=> West | 131 (141) | [$]< 0.0001 | 40 (35) | [$]0.011 | 91 (106) | [$]<0.0001 | 35 (28) | [$]<0.0001 | 96 (113) | [$]0.0021 |
| [%€]East <=> West | 263 (203) | [$]0.0014 [£]0.0083 | 44 (38) | [$]<0.0001 [£]0.0047 | 219 (165) | [$]0.8 [£]<0.0001 | 48 (47) | [$]<0.0001 [£]0.62 | 215 (156) | [$]0.79 [£]0.0007 |
| Intra-farm East | 49 (65) | | | | | | | | | |
| Intra-farm West | 34 (41) | [*]0.4566 | | | | | | | | |

[#]Number of isolate pairs with less than 21 different alleles by cgMLST and in parenthesis, numbers of isolates with at least one match. [§]Two-tailed exact Fisher test on the proportion of isolates with at least one match. [@]At least one of the two isolates showed an AmpC-like phenotype. [€]Excluding intra-farm pairs. [%]For isolates pairs on both sides of the Meuse, the average of the number of isolates in each compartment was used for statistical analyses. [$]Compared with East-East. [£]compared with West-West. *Compared with Intra-farm East.

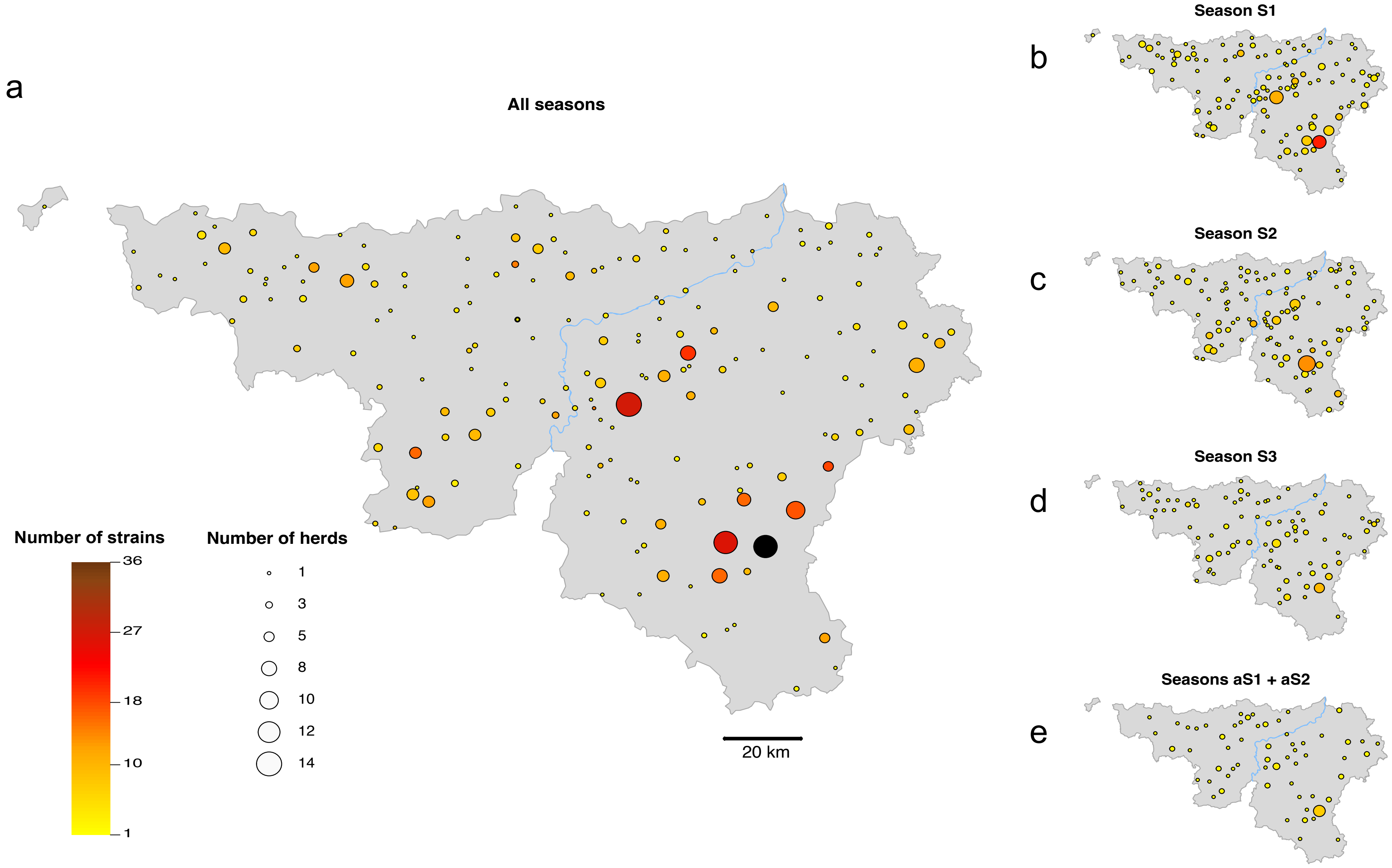


**Fig. S1. Geographic distribution of farms and animals across the Wallonia.** Number of farms and animals in each postal code providing samples are represented by colored disk. Disk were positioned according to the geographic coordinates of the postal code (from the Open Data Wallonie-Bruxelles web site https://www.odwb.be/). The size of the disk and its color indicate the number of herds and the number of *E. coli* isolates in each postal code, respectively, according to the figure key. **a** isolates from the five calving seasons, **b** season S1, 2017-2018, **c** season S2, 2018-2019, **d** season S3, 2019-2020, **e** season aS1 and aS2, 2015-2017.

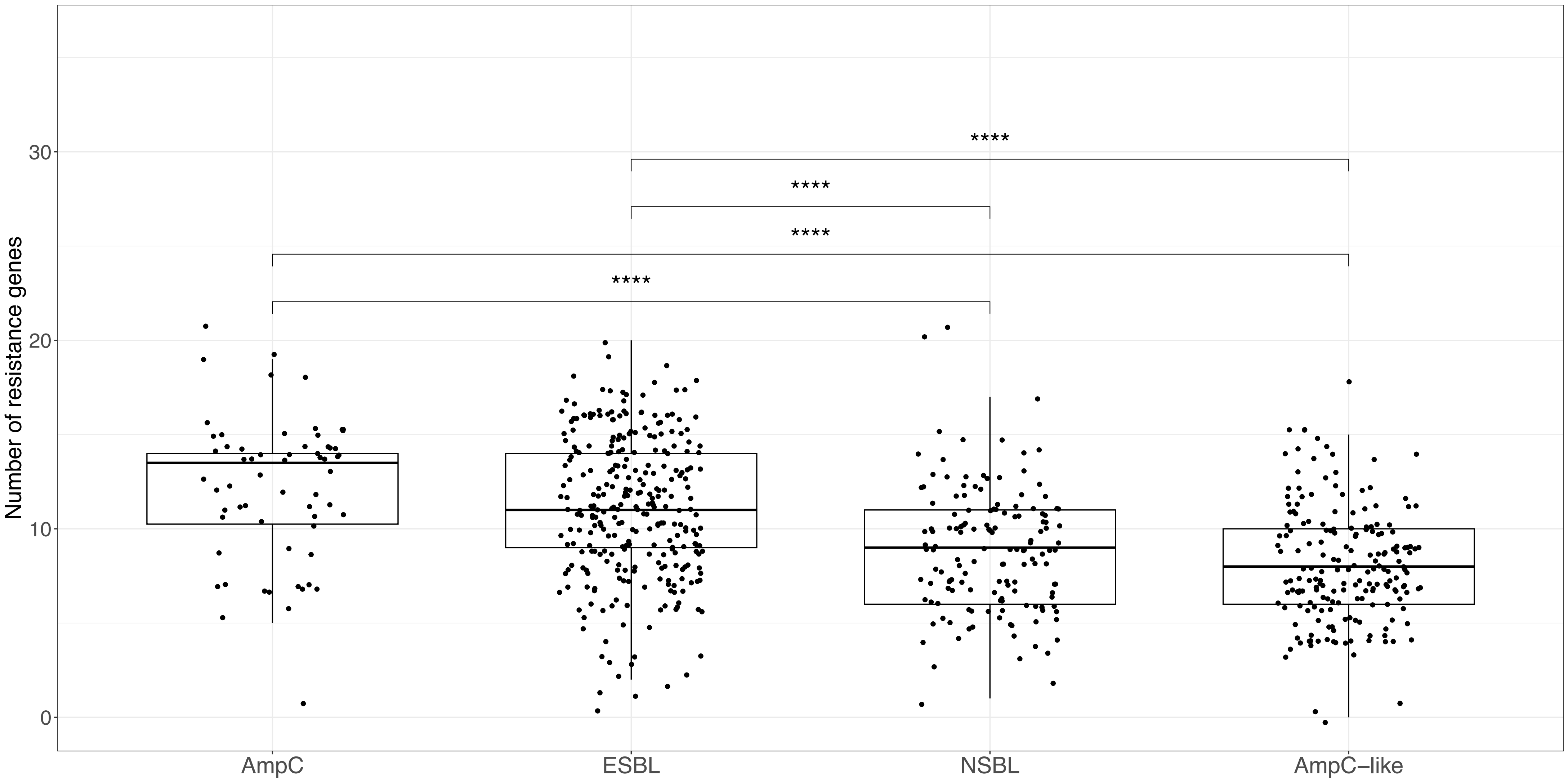


**Fig S2**: **Number of acquired resistance genes per isolate according to the ß-lactam resistance profile.** Isolates were from the three calving seasons S1-3. The horizontal lines in the boxes represent the median number of ARGs. The box boundaries represent the first and third quartiles of the distribution and box-plot whiskers span 1.5 times the interquartile range of the distribution. Group differences were first assessed using a global Kruskal-Wallis test, followed by pairwise Dunn's post-hoc tests with Benjamini-Hochberg correction for multiple comparisons. Significant differences were observed between most groups, **** p-value < 0.00005

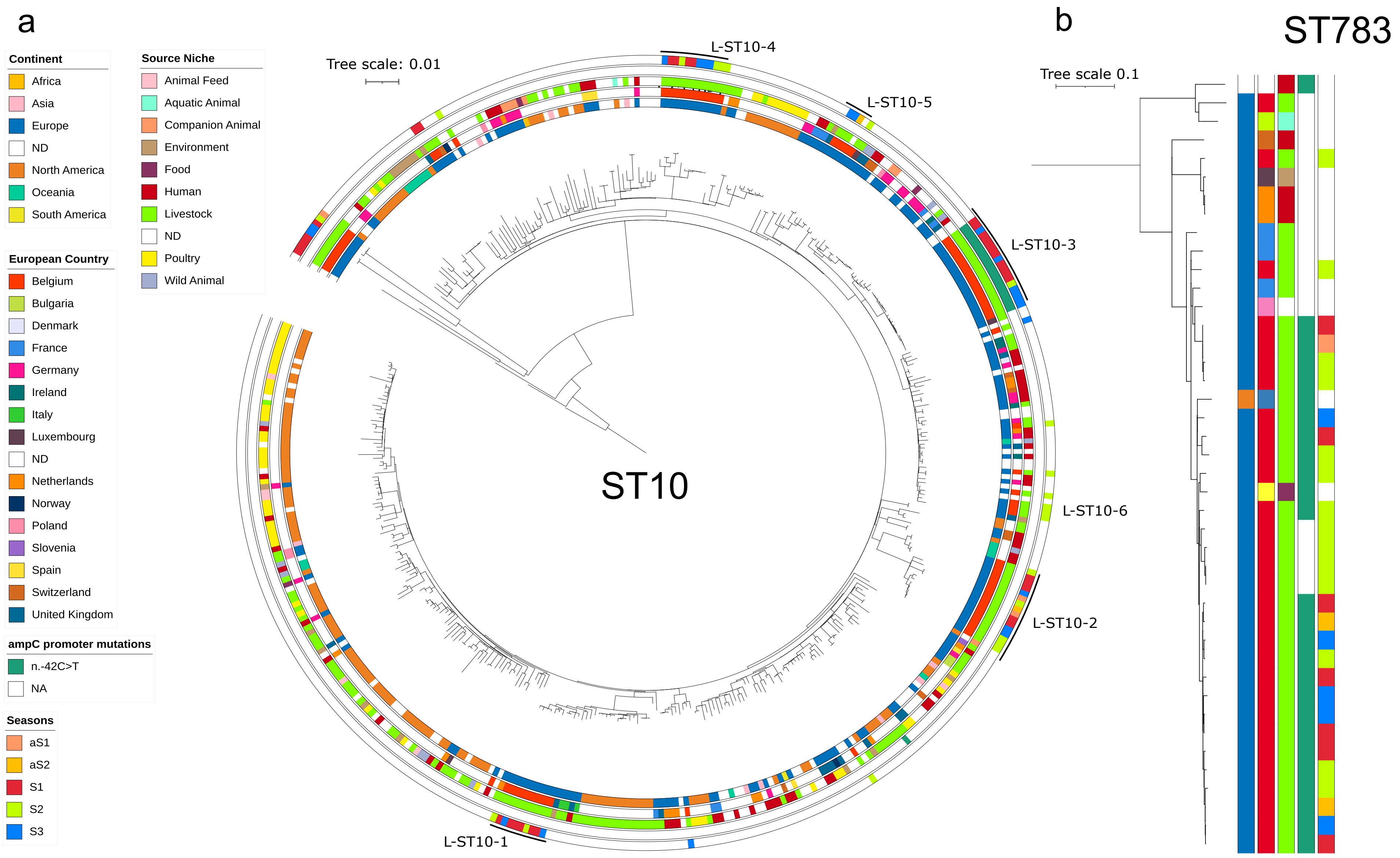


**Fig. S3 Phylogeny and characteristics of ST10 and ST783 *E. coli* isolates from the five calving seasons combined with EnteroBase isolates.** Maximum likelihood phylogeny was performed by using RAxML 8.2.12 [1] based on recombination-free core genome alignment, computed with Snippy 4.6.0 (https://github.com/tseemann/snippy) and Gubbins 2.4.1 to filter recombined regions [2]. **a.** ST10. The tree was built by including 353 ST10 isolates from EnteroBase with a cgMLST distance of less than 100 alleles with isolate R0001 which belong to the lineage L-ST10-3. Among the 267 known hosts, 71 were bovine, 63 human et 40 poultry. **b**. ST783. Only 13 isolates were retrieved from EnteroBase including four of human origin. Genomes were annotated according to the figure key as circles from inside to outside (A) or from left to right (B): the continent of origin, country, source niche, $P_{ampC}$ mutation and the season. The cluster reported in table 2 are indicated outside the last circle (A).

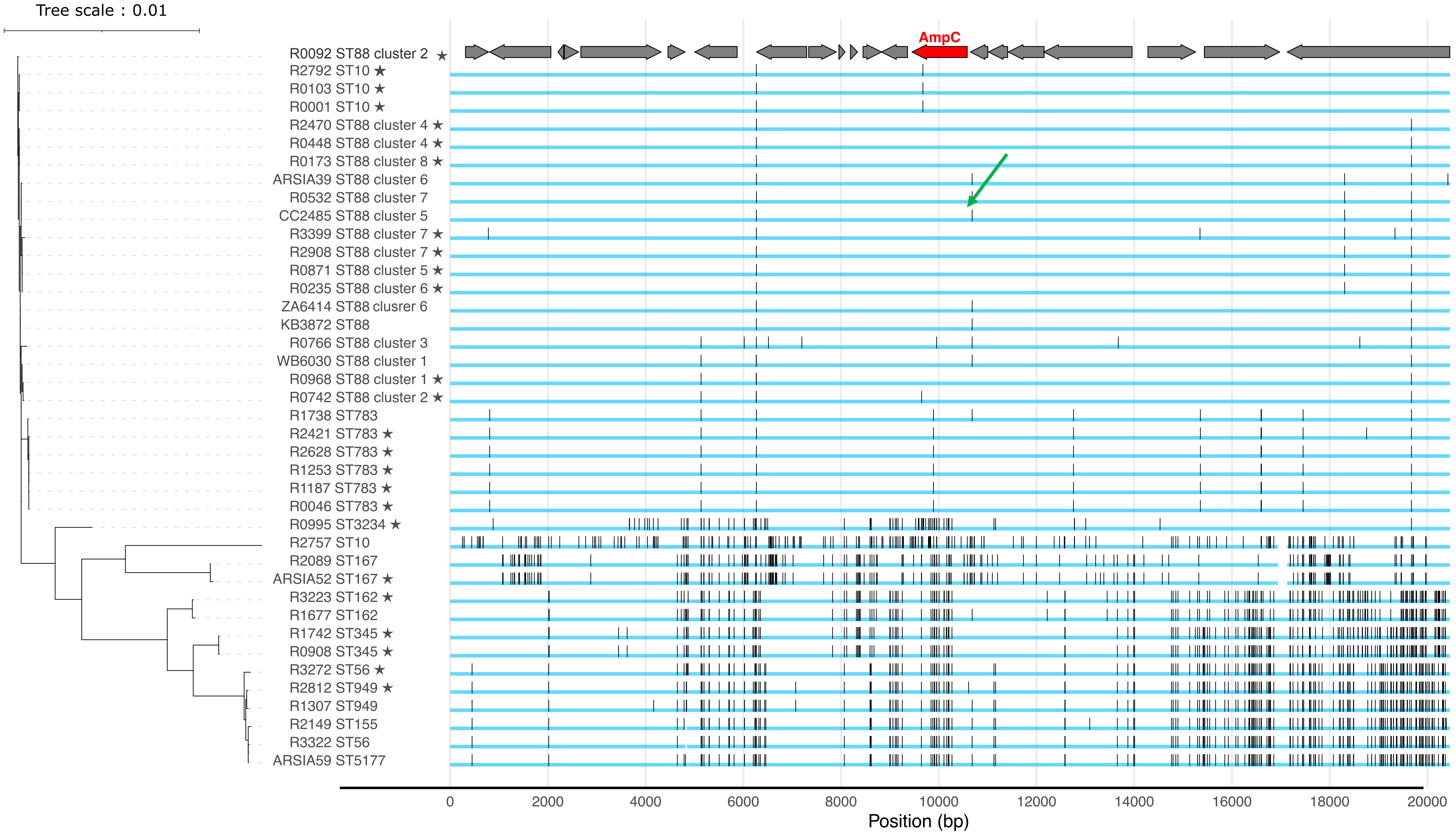


**Fig. S4. Recombination detection at the *ampC* locus**. SNPs in the chromosomal region 10 kb upstream and downstream *ampC* in 23 isolates mutated in $P_{ampC}$ (-42) and in 15 closely related isolates non mutated. SNP were identified by using Breseq [3]. Reference sequence was R0092 from lineage L-ST88-2 and mutated in $P_{ampC}$. SNPs are indicated by small vertical lines. Recombination was detected only in the three isolates belonging to the lineage L-ST10-3. The $P_{ampC}$ variant (wild type) is highlighted by a green arrow. The *ampC* gene is shown by a red arrow. Sequences were ordered according to the phylogeny of the visualized *ampC* region shown on the left part.

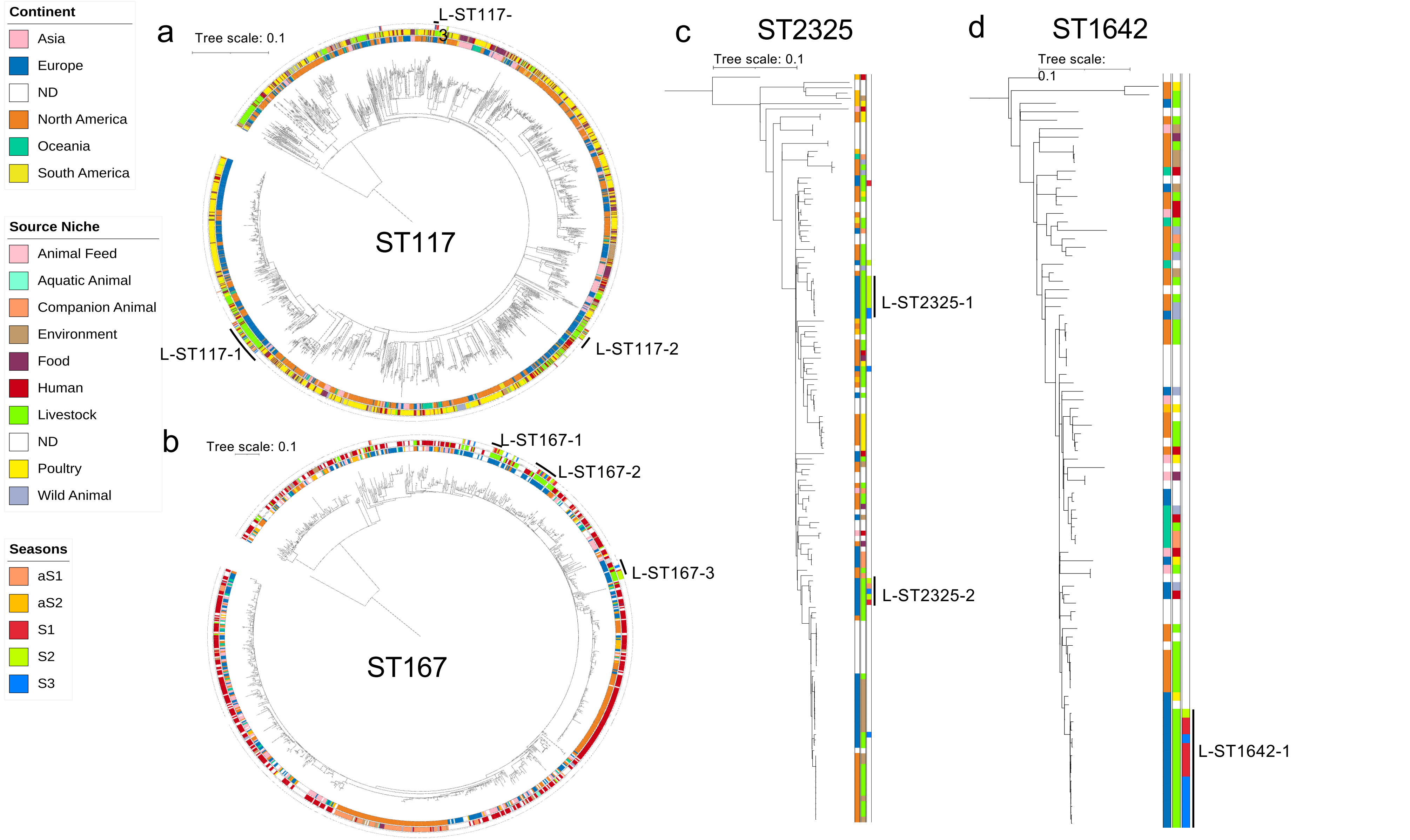


**Fig. S5. Phylogeny and characteristics of ST117, ST167, ST2325 and ST1642 *E. coli* from the five seasons combined with EnteroBase isolates**. Maximum likelihood phylogeny was performed by using RAxML 8.2.12 [1] based on recombination-free core genome alignment, computed with Snippy 4.6.0 and Gubbins 2.4.1 [2] to filter recombined regions. **a**. ST117 (n=71). the tree was built by including only EnteroBase sequences with country and host information (n=1525). 67% of the isolates were from poultry origin. **b**. ST167 (n=44). The tree was built by including 989 ST167 isolates from EnteroBase. Among the 734 with known hosts, 532 were of human origin and 25 of bovine origin. **c**. ST2325 (n=17). 123 isolates were retrieved from EnteroBase, with five of human origin and 22 of bovine origin from the 90 with known host. **d**. ST1642 (n=14). 81 isolates were retrieved from EnteroBase (57 annotated), with seven of human origin and 13 of bovine origin. Genomes were annotated according to the figure key as circles from inside to outside (a and b) or from left to right (c and d): continent of origin, source niche, and season of isolation for the isolates of the study. The cluster reported in table 2 are indicated outside the last circle (a and b) or on the right of the tree (c and d).

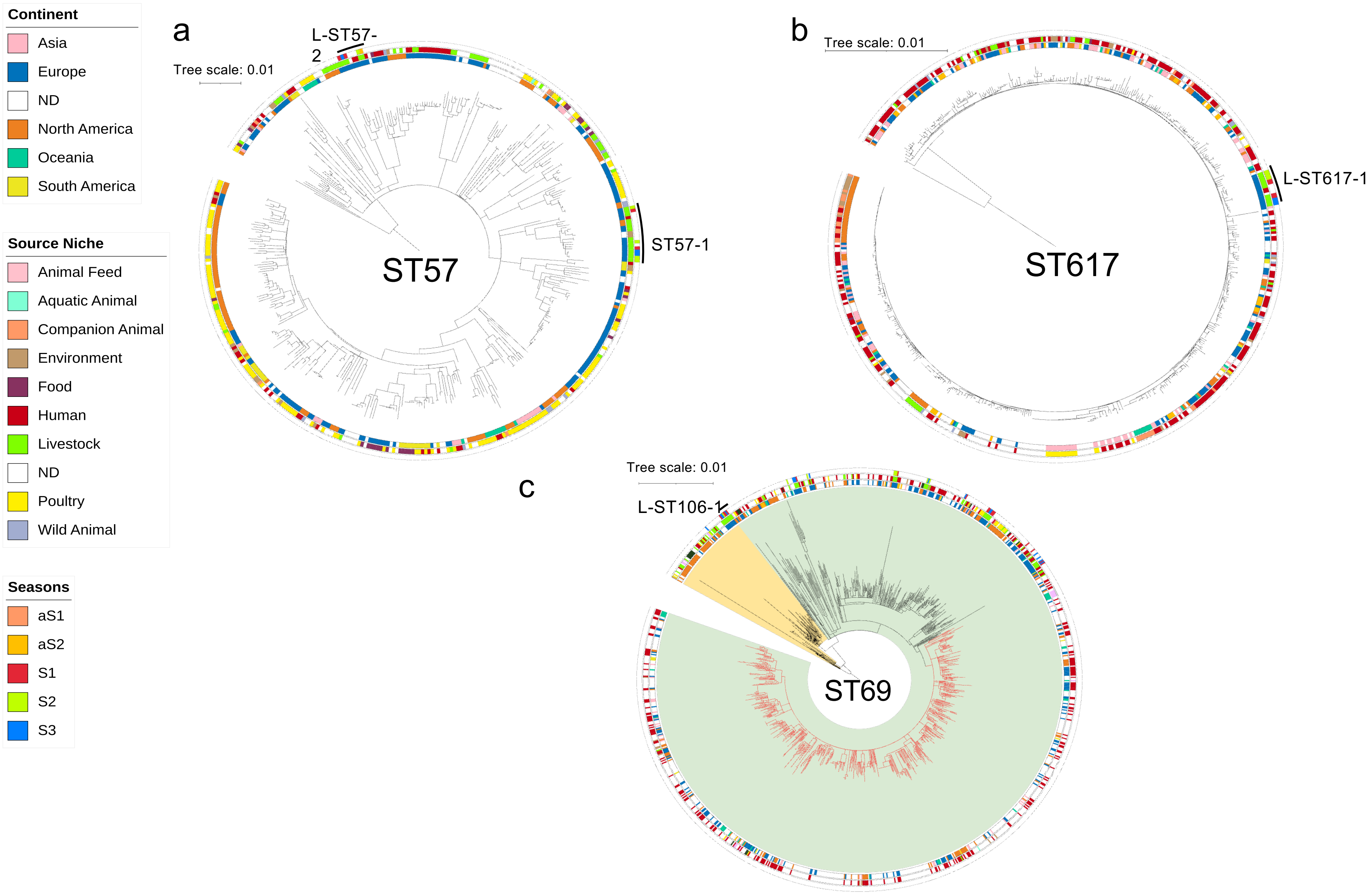


**Fig. S6. Phylogeny and characteristics of ST57, ST617 and ST69/ST106 *E. coli* isolates from the five seasons combined with EnteroBase isolates.** Maximum likelihood phylogeny was performed by using RAxML 8.2.12 [1] based on recombination-free core genome alignment, computed with Snippy 4.6.0 and Gubbins 2.4.1 [2] to filter recombined regions. **a.** ST57, 394 EnteroBase isolates (311 annotated), 50 of human and 20 of bovine origin. **b**, ST617, 561 EnteroBase isolates (386 annotated), 256 of human and 15 of bovine origin. **c**, ST69/ST106, 1103 EnteroBase isolates (467 annotated), 302 of human and 68 of bovine origin. In ST69, note that no bovine isolate cluster with the late appearing clade gathering only human isolates (in red). Genomes were annotated according to the figure key as circles from inside to outside: the continent of origin, the source niche, and the season of isolation. The cluster reported in table 2 are indicated outside the last circle.

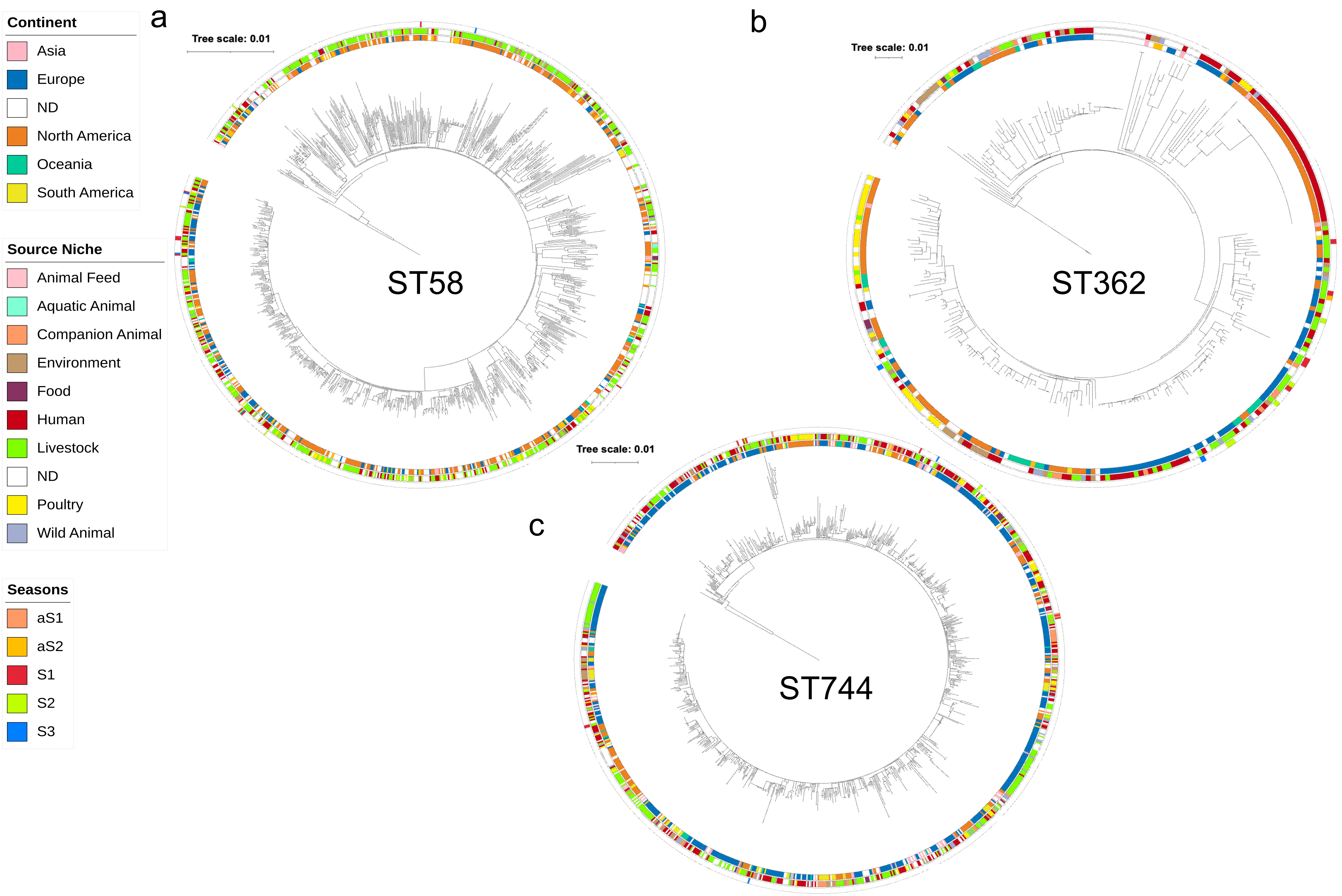


**Fig. S7. Phylogeny and characteristics of ST58, ST362 and ST744 *E. coli* isolates from the five seasons combined with EnteroBase isolates.** Maximum likelihood phylogeny was performed by using RAxML 8.2.12 [1] based on recombination-free core genome alignment, computed with Snippy 4.6.0 and Gubbins 2.4.1 [2] to filter recombined regions. **a.** ST58 2730 EnteroBase isolates (2140 with sources), 360 of human and 536 of bovine origin. **b**, ST362, 310 EnteroBase isolates (248 annotated), 100 of human and 34 of bovine origin. **c**, ST744, 1012 EnteroBase isolates (831 annotated), 260 of human and 49 of bovine origin. Genomes were annotated according to the figure key as circles from inside to outside: the continent of origin, the source niche, and the season of isolation. No cluster was identified in these three STs

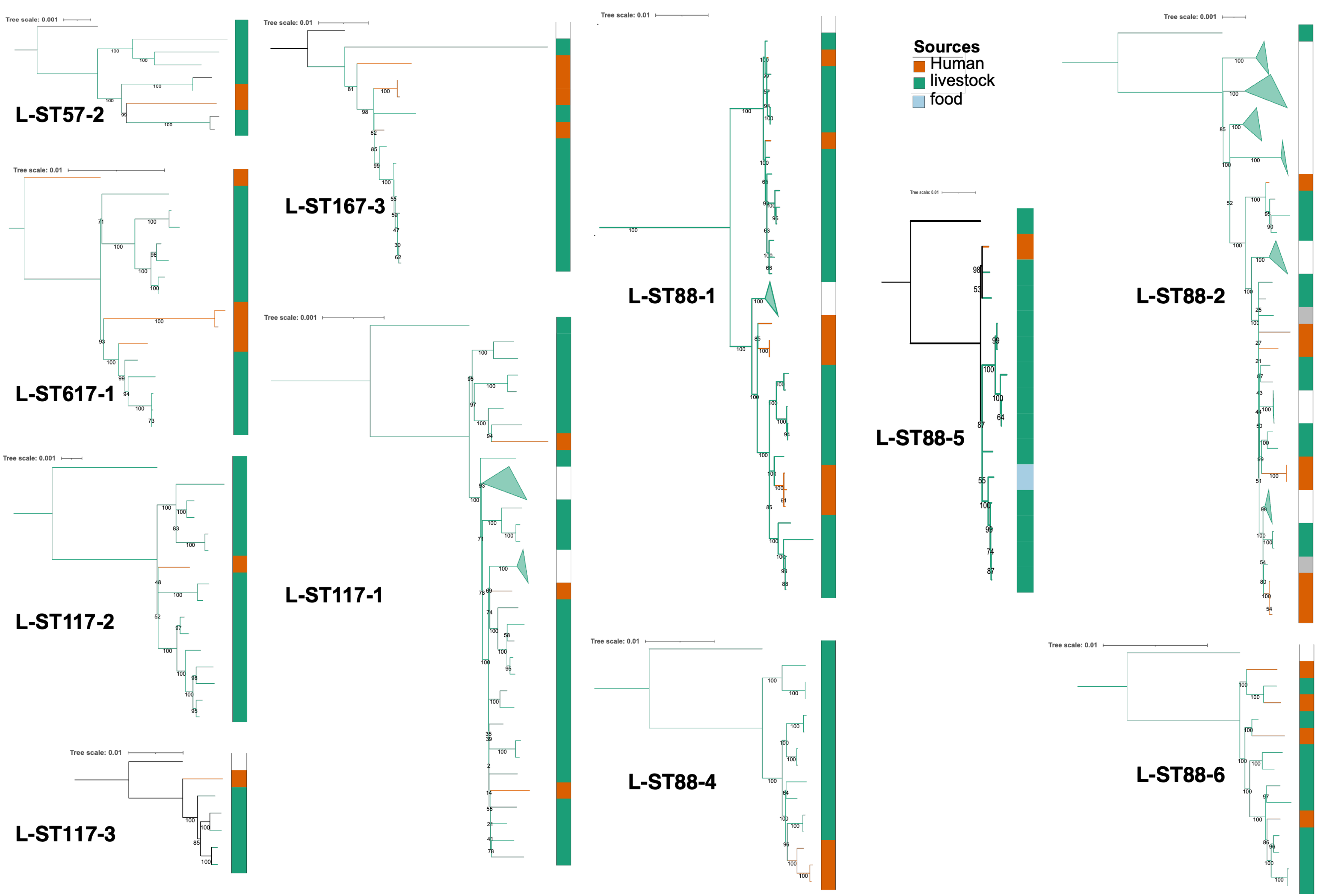


**Fig. S8. Ancestral host of the 11 lineages including EnteroBase isolates of human origin**. Phylogenetic trees and PastML [4] host predictions were performed on all the isolates and nodes of an ST and sub-trees and predictions of each lineage were extracted from these trees. Host probabilities of the ancestral nodes are indicated in green, bovine more likely, or in red, human more likely. Bovine isolates are in green and human isolates in red. Bootstrap values are in black. Details on lineages are in Table 2.

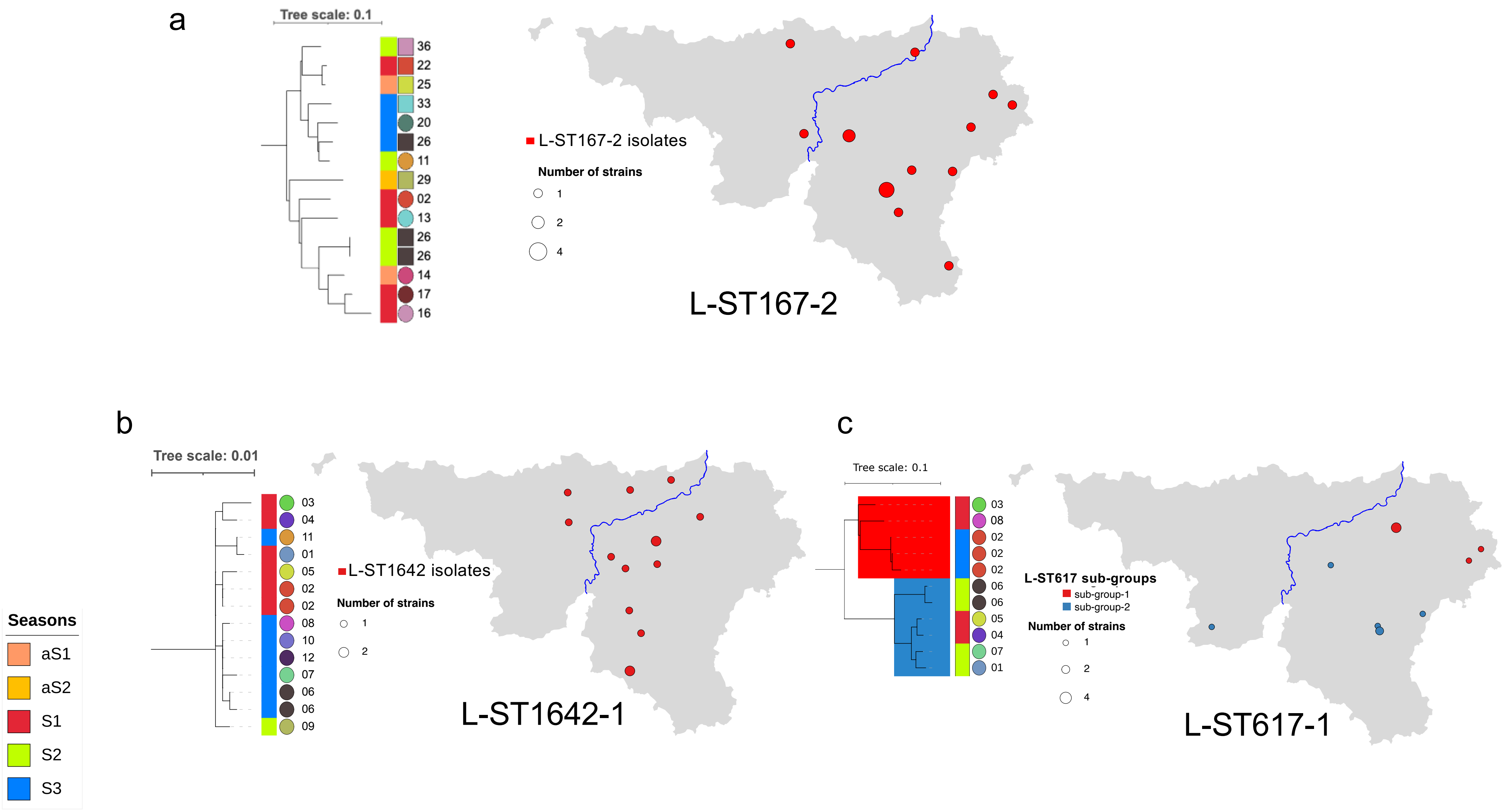


**Fig. S9. Local dissemination of closely related isolates (ST167, ST1642, ST617) and geographic specificities.** The geographic localization was compared to the phylogenetic proximity by localizing isolates according to the GPS coordinates of the postal code. Disk diameters correspond to the number of isolates. Colors correspond to the branch on the tree. Isolates were annotated on the right of the tree according to the figure key. From left to right, the season (in white, EnteroBase isolates) and the herd. The blue line represents the Meuse. **a.** L-ST167-2, isolates were mainly on the east, **b.** L-ST1642-1, **c.** L-ST617-1

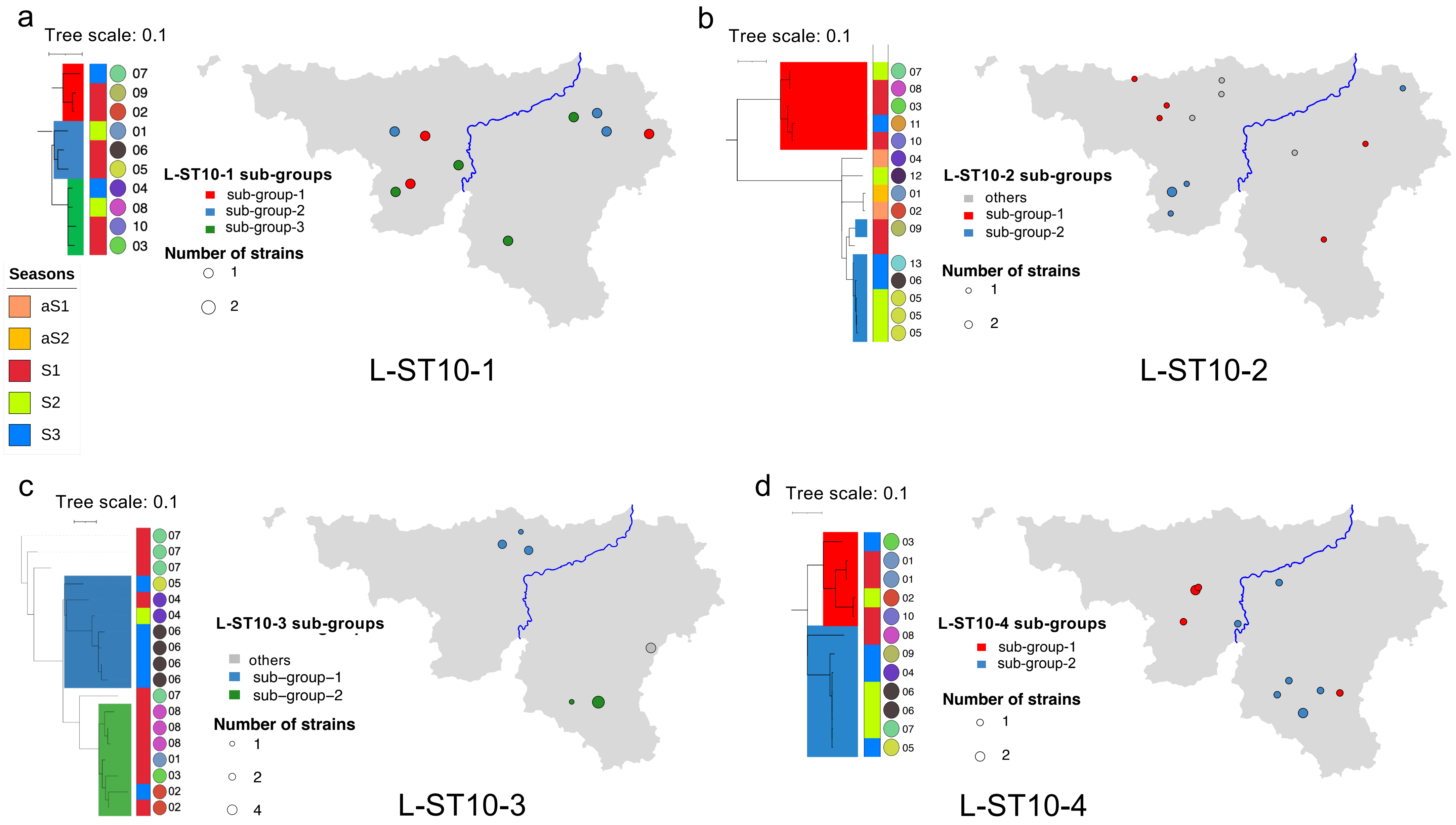


**Fig. S10. Local dissemination of closely related isolates (ST10) and geographical specificities.** The geographic localization was compared to the phylogenetic proximity by localizing isolates according to the GPS coordinates of the postal code. Disk diameters correspond to the number of isolates. Colors correspond to the branch on the tree. Isolates were annotated on the right of the tree according to the figure key. From left to right, the season (in white, EnteroBase isolates) and the herd. The blue line represents the Meuse. **a.** L-ST10-1, **b.** L-ST10-2, **c.** Lineage L-ST10-3, **d.** Lineage L-ST10-4.

### References


1. Stamatakis A. Raxml version 8: A tool for phylogenetic analysis and post-analysis of large phylogenies. *Bioinformatics*. 2014;**30**:1312-3 https://doi.org/10.1093/bioinformatics/btu033
2. Croucher NJ, Page AJ, Connor TR *et al.* Rapid phylogenetic analysis of large samples of recombinant bacterial whole genome sequences using gubbins. *Nucleic Acids Res*. 2015;**43**:e15 https://doi.org/10.1093/nar/gku1196
3. Deatherage DE, Barrick JE. Identification of mutations in laboratory-evolved microbes from next-generation sequencing data using BRESEQ. *Methods Mol Biol*. 2014;**1151**:165-88 https://doi.org/10.1007/978-1-4939-0554-6_12
4. Ishikawa SA, Zhukova A, Iwasaki W *et al.* A fast likelihood method to reconstruct and visualize ancestral scenarios. *Mol Biol Evol*. 2019;**36**:2069-85 https://doi.org/10.1093/molbev/msz131
